# PU.1 inhibition sensitizes stem-monocytic AML to BCL2 blockade

**DOI:** 10.64898/2026.01.20.700677

**Authors:** William M Yashar, Itallia V Pacentine, Akram Taherinasab, Thai Nguyen, Samantha Worme, Mitsuhiro Tsuchiya, Theresa Lusardi, Chelsea Hutchinson, Thomas L Fillmore, Camilo Posso, Sara JC Gosline, Paul D Piehowski, Galip Gurkan Yardimci, Andrew C Adey, Julia E Maxson, Theodore P Braun

**Affiliations:** Division of Oncologic Sciences, Department of Medicine, Oregon Health & Science University; Portland, OR, 97239, USA; Knight Cancer Institute, Oregon Health & Science University; Portland, OR, 97239, USA; Cancer Early Detection Advanced Research Center, Oregon Health & Science University; Portland, OR, 97239, USA; Biological Sciences Division, Pacific Northwest National Laboratory, Richland WA; Environmental and Molecular Sciences Division, Pacific Northwest National Laboratory, Richland WA; Department of Molecular and Medical Genetics, Oregon Health and Science University; Portland, OR 97239, USA; Division of Hematology & Medical Oncology, Department of Medicine, Oregon Health & Science University; Portland, OR, 97239, USA; Cell Developmental & Cancer Biology, Oregon Health & Science University; Portland, OR, 97239, USA

**Keywords:** Venetoclax, BCL2 inhibition, PU.1, myeloid differentiation, stem-monocytic acute myeloid leukemia

## Abstract

Acute myeloid leukemia (AML) exhibits substantial transcriptional heterogeneity across differentiation states that influences therapeutic response to BCL2 inhibition with venetoclax. While hematopoietic stem cell (HSC)-like AMLs show high sensitivity to venetoclax and monocytic-like AMLs demonstrate resistance, the therapeutic behavior of leukemias harboring both transcriptional programs remains poorly defined. Analysis of a large AML cohort reveals a distinct patient population exhibiting concurrent HSC- and monocyte-like transcriptional signatures, which we term stem-monocytic AML. *Ex vivo* drug sensitivity profiling demonstrates that stem-monocytic AMLs exhibit venetoclax resistance comparable to pure monocytic disease, despite expressing HSC-like transcriptional features. Using a leukemia cell line model that recapitulates stem-monocytic AML characteristics, we show through immunophenotyping and single-cell lineage tracing that venetoclax preferentially depletes immature blasts while sparing differentiated monocytic populations. Single-cell transcriptomic and chromatin accessibility analyses identify enrichment of myeloid differentiation transcription factors, particularly PU.1, in resistant populations. A targeted CRISPR knockout screen confirms that PU.1 disruption induces differentiation arrest and enhances venetoclax sensitivity primarily in the immature immunophenotypic compartments. Pharmacologic PU.1 inhibition with the small molecule DB2313 synergizes with venetoclax in both cell line models and primary patient samples. These findings establish stem-monocytic AML as a transcriptionally and functionally distinct subtype and nominate combined PU.1 and BCL2 inhibition as a rational therapeutic strategy for improving venetoclax response in this patient population.

## INTRODUCTION

Acute myeloid leukemia (AML) is an often-fatal hematologic malignancy characterized by uncontrolled proliferation of immature myeloid blasts. Although AML arises from arrested myeloid differentiation, leukemic blasts exhibit morphologic and immunophenotypic features spanning a range of maturation stages in myeloid development. The French–American–British classification codified these distinctions by categorizing AML subtypes according to morphologic differentiation^1,2^. However, molecular and cytogenetic classifications have largely supplanted these subtypes, offering superior prognostic and therapeutic prediction^3,4^. Despite this shift toward genetic characterization, the focus on differentiation heterogeneity in AML has reemerged with transcriptional profiling. Single-cell RNA sequencing (scRNA-seq) studies reveal that leukemic blasts occupy a continuous spectrum of differentiation states, ranging from immature hematopoietic stem cell-like to mature monocyte-like program^5,6^. Quantitative scoring systems now capture this gradient, linking differentiation state transcriptional signatures to clinical and biological behavior^7,8^. This recognition has renewed interest in how leukemic differentiation state shapes the therapeutic behavior of AML, particularly in response to the BCL2 inhibitor, venetoclax.

The differentiation state of AML blasts is increasingly recognized as a major determinant of therapeutic response. Venetoclax has shown marked efficacy in transcriptionally immature AMLs^9–11^. Recent analysis of a large international cohort has demonstrated that monocytic AML is associated with decreased overall survival in the setting of venetoclax-based therapy, further validating leukemic differentiation state as a functional biomarker of response^12^. Most investigations, however, dichotomize AMLs into immature or mature states, neglecting leukemias composed of both hematopoietic stem cell (HSC)-like and monocytic-like transcriptional programs. These mixed transcriptional states complicate current classification schemes and raise key questions regarding how differentiation heterogeneity influences treatment resistance.

We sought to address this gap by examining AMLs exhibiting concurrent HSC- and monocytic-like transcriptional features, which we term stem-monocytic AMLs. Using a large patient cohort, we found that while stem-monocytic AMLs are not identifiable by conventional mutational or surface marker profiling, they represent a unique AML subtype with a venetoclax resistance profile comparable to that of monocytic AMLs. These findings demonstrate that differentiated leukemic blasts confer resistance to venetoclax even in the presence of immature disease. To study the relationship between differentiation state and venetoclax sensitivity, we identified a leukemic stem cell line that maintains a differentiation spectrum and recapitulates maturation state-dependent responses to venetoclax^13^. In this cell line, we used scRNA-seq and single-cell ATAC sequencing (scATAC-seq) to identify key factors involved in venetoclax resistance. Guided by these findings, we performed a CRISPR knockout screen that identified inhibition of PU.1, a myeloid pioneer transcription factor, as a rational combination partner with venetoclax. Pharmacologic inhibition of PU.1 enhanced venetoclax efficacy in stem-monocytic AML cell line models and in primary patient samples, revealing a tractable therapeutic vulnerability in this population^14,15^. These findings define stem-monocytic AML as a distinct transcriptional and functional subtype and highlight the targeting of lineage-defining transcriptional programs to enhance sensitivity to BCL2 inhibition in this patient population.

## MATERIALS AND METHODS

### Cell culture

#### General

All cells were cultured at 5% CO₂ and 37°C. Cell lines were tested for mycoplasma concentration at the time of freezing as well as monthly for any cell lines in culture.

#### OCI-AML8227

OCI-AML8227 cells were provided by John Dick (University of Toronto) under a materials transfer agreement. These cells were shipped frozen on dry ice and then thawed in a water bath set to 37°C. Thaw media consisting of 49.5% FBS (HyClone), 49.5% X-VIVO 10 (Lonza #04-380Q), and 1% DNAse I (Roche #11284932001) was slowly added drop-wise to the cells. They were incubated for 10 minutes at room temperature and then washed twice with DPBS (Gibco). After washing, the cells were transferred into culture media consisting of sterile filtered (0.22 μm, EMD Millipore #SE1M179M6) 20% BIT (StemCell Technologies #9500) and 80% X-VIVO 10 supplemented with IL-3 (10 ng/mL; Peprotech #200-03), SCF (100 ng/mL; Peprotech #300-07), IL-6 (20 ng/mL; Peprotech #200-06), Flt3-Ligand (50 ng/mL; Peprotech #300-19), G-CSF (10 ng/mL; Peprotech #300-23), and TPO (25 ng/mL; Peprotech #300-18). Unless otherwise stated, OCI-AML8227 cells were cultured in the inner eight wells of a 24-well plate with the outer wells filled with DPBS. Additionally, OCI-AML8227 cells were seeded at a concentration of 4e5 cells per well and then incubated for 48 hours before experiments.

#### Other cell lines

All other cell lines were cultured in standard incubation conditions according to the table below:

| Cell line | Growth media |
| --- | --- |
| MUTZ3 | Use ¼ old media + ¾ fresh (60% α-MEM* + 20% FBS<sup>†</sup> + 20% 5637 media) |
| U937 | RPMI<sup>§</sup> + 10% FBS<sup>†</sup> |
| MOLM13 | RPMI<sup>§</sup> + 10% FBS<sup>†</sup> |
| KG-1a | 80% α-MEM* + 20% FBS<sup>†</sup> |
| AML3 | 80% α-MEM* + 20% FBS<sup>†</sup> |
| THP1 | RPMI<sup>§</sup> + 10% FBS<sup>†</sup> |
| 5637 | RPMI<sup>§</sup> + 10% FBS<sup>†</sup> |

\*α-MEM = MEM Alpha Modification (1X) + L-Glutamine + Ribonucleosides and Deoxyribonucleosides (HyClone # SH30265.01),
<sup>†</sup> FBS = fetal bovine serum (HyClone # SH3039603), heat-inactivated at 55°C for 30 minutes
<sup>§</sup> RPMI = RPMI Medium 1640 (1X) (gibco # 11875-093)

#### Patient samples

All patients gave written informed consent to participate in this study, which was conducted in accordance with the Declaration of Helsinki and had the approval and guidance of the institutional review boards at Oregon Health & Science University (OHSU; IRB#9570; #4422; NCT01728402). Samples were sent to the coordinating center where they were coded and processed. Mononuclear cells were isolated by Ficoll gradient centrifugation from freshly obtained bone marrow aspirates or peripheral blood draws. Clinical, prognostic, genetic, cytogenetic, and pathologic lab values as well as treatment and outcome data were manually curated from patient electronic medical records. Genetic characterization of the leukemia samples included results of a clinical deep-sequencing panel of genes commonly mutated in hematologic malignancies (GeneTrails, OHSU Knight Diagnostics Laboratory).

#### Immunomagnetic fractionation

OCI-AML8227 cells were immunomagnetically fractionated for CD34 surface expression using CD34 Microbeads (Miltenyi Biotec #130-046-702). Cells were resuspended in sterile filtered (0.22 μm) MACS buffer (0.5% BSA and 2 mM EDTA). Cells were incubated with CD34 Microbeads for 30 minutes at 4°C. The beads were washed from the media, resuspended in MACS buffer, and then passed through a MACS Cell Separator LS column (Miltenyi Biotec #130-042-401) with a pre-separation cell strainer (Miltenyi Biotec #130-101-812). The column was washed three times with MACS buffer. CD34-positive OCI-AML8227 cells were retained in the column and eluted after removal from the magnetic field. CD34-negative cells were isolated from the LS column flow-through.

#### Flow cytometry

Cultured cells were first counted using a TC20 Automated Cell Counter (Bio-Rad #1450102). After counting, cells were washed with 1x DPBS and incubated with 1:100 FcR Blocking Reagent (Miltenyi Biotec #130-046-702) and 1:1500 Live/Dead Blue (Invitrogen #L23105) at room temperature for 20 minutes. Cells were then washed with Stain Buffer (BD Biosciences #554656) and stained with the following antibodies at 4°C for 30 minutes: CD14-BV421 (BioLegend #301830), CD64-FITC (BioLegend #305006), CD34-PE (BD Biosciences #348057), and CD38-APC-Fire810 (BioLegend #356644). After staining, cells were washed with Stain Buffer and analyzed using a BD Symphony A5 flow cytometer.

#### Phosphoproteomics

##### Sample preparation

Cultured, immunomagnetically fractionated OCI-AML8227 cells were plated at a concentration of 1e6 cells per well in 2.5 mL media. Three replicates per condition were performed. Cells were allowed to recover and expand for 2 days, then treated with either DMSO or 1 μM venetoclax (MedChem Express #HY-15531) and collected at 0 hour (untreated), 30 minutes, and 6 hours post-treatment. The entire well contents were collected and centrifuged at 500g for 3 minutes in a swing bucket rotor, washed with DPBS, centrifuged again to pellet, supernatant aspirated, and snap frozen in liquid nitrogen before storing at -80°C. Samples were shipped overnight on dry ice to the Pacific Northwest National Laboratory for mass spectrometry analysis.

##### Protein digestion and labeling

Sample preparation for proteomics followed the protocol developed under the CPTAC consortium^16^. PBS washed cell pellets were lysed with 100 μL fresh lysis buffer, 8 M urea (Sigma-Aldrich), 50 mM Tris pH 8.0, 75 mM sodium chloride, 1 mM ethylenediamine tetra-acetic acid, 2 μg/mL Aprotinin (Sigma-Aldrich), 10 μg/mL Leupeptin (Roche), 1 mM PMSF in EtOH, 10 mM sodium fluoride, 1% of phosphatase inhibitor cocktail 2 and 3 (Sigma-Aldrich), 20 μM PUGNAc, and 0.01 U/μL Benzonase. Lysates were vortexed for 10 seconds and then placed in thermomixer set for 15 minutes at 4°C and 800 RPM, vortexing was repeated and samples were incubated again for 15 minutes with the same settings. Samples were centrifuged for 10 minutes at 4°C and 18000 rcf to remove cell debris. The supernatant was removed and transferred to a fresh tube. A BCA (ThermoFisher) assay was used on the supernatant to measure the protein yield.

Samples were all diluted to 2 µg/µL protein concentration with fresh lysis buffer prior to starting digestion. The samples were reduced with 5 mM dithiothreitol (DTT; Sigma-Aldrich) for 1 hour at 37°C and 800 rpm. Reduced cysteines were alkylated with 10 mM iodacetamide (IAA; Sigma-Aldrich) for 45 minutes at 25°C and 800 rpm in the dark. The samples were diluted 4-fold with 50 mM Tris HCL pH 8.0 and then Lys-C (Wako) was added at a 1:20 enzyme:substrate ratio, followed by incubation for 2 hours at 25°C, shaking at 800 rpm. Trypsin (Promega) was then added at a 1:20 enzyme:substrate ratio, followed by a 14-hour incubation at 25°C and and shaking at 800 rpm. The digestion was quenched with formic acid at 1% by volume followed by centrifugation for 15 minutes at 1500 rcf, to remove any remaining cell debris. Peptides samples were desalted using a C18 solid phase extraction (SPE) cartridge (Waters Sep-Pak).

After SPE, samples were dried down and reconstituted with 50 mM HEPES, pH 8.5 to a concentration of 5 μg/μL. Tandem mass tag (TMT) reagent aliquots were dissolved in 250 μL of anhydrous acetonitrile to a final concentration of 20 μg/μL. The tag was added to the sample at a 1:1 peptide:label ratio and incubated for 1 hour at 25°C and 400 rpm and then diluted to 2.5 mg/mL with 50 mM HEPES pH 8.5, 20% acetonitrile (ACN). The labeling reaction was quenched with 5% hydroxylamine and incubated for 15 minutes at 25°C and 400 rpm. The samples were then combined to create two plex sets and concentrated in a speed-vac before a final C18 SPE cleanup. Each 16-plex experiment was fractionated into 96 fractions by high pH reversed phase separation, followed by concatenation into 12 global fractions for MS analysis.

##### Phosphopeptide enrichment using IMAC

Six samples per plex were then created by concatenation from the 12 global TMT fractions. Fe^3+^-NTA-agarose beads were freshly prepared for phosphopeptide enrichment using Ni-NTA-agarose beads (Qiagen). Sample peptides were reconstituted to a 0.5 μg/μL concentration with 80% ACN, 0.1% TFA and incubated with 40 μL of the bead suspension for 30 minutes at RT in a thermomixer set at 800 rpm. After incubation, the beads were washed with 100 μL 80% ACN, 0.1% TFA and 50 μL 1% FA to remove any non-specific binding. Phosphopetides were eluted off beads with 210 μL 500 mM K_2_HPO_4_, pH 7.0 directly onto C18 stage tips and eluted from C18 material with 60 μL 50% ACN, 0.1% FA. Samples were dried in speed-vac concentrator and reconstituted with 12 μL 3% ACN, 0.1% FA.

##### LC-MS/MS Analysis

Sample fractions were separated on a Waters nano-Aquity UPLC system (Waters) equipped with a homemade 75 um I.D. x 25 cm length C18 column packed with 1.7 um BEH C18 particles (Waters). A 120-minute gradient of 95% mobile phase A (0.1% (v/v) formic acid in water) to 19% mobile phase B (0.1% (v/v) FA in acetonitrile) was applied to each fraction. The separation was coupled to a Thermo Orbitrap Exploris mass spectrometer. MS Spectra were collected from 350 to 1800 m/z at an MS1 resolution of 60K. A top 20 method was used for the collection of MS2 spectra at a mass resolution of 45K. An isolation window of 0.7 m/z was used for higher energy collision dissociation (HCD), singly charged species were excluded, and the dynamic exclusion window was 30 seconds.

#### Single-cell barcoding

##### Library preparation

Lentiviral barcoding was performed using the pLARRY-EGFP-BCv1 plasmid as previously described^17^. pLARRY-EGFP-BCv1 was a gift from Fernando Camargo (Addgene plasmid #140025). The pLARRY plasmid was retransformed into STBL4 ElectroMAX competent cells (ThermoFisher #11635018) and plated onto large LB-agar plates supplemented with 100 μg/mL ampicillin. Library plasmid DNA was purified using the NucleoBond Xtra Maxi EF kit (Macherey-Nagel #740424.50). Bacterial cultures were extracted using the DNeasy Blood & Tissue Kit (Qiagen #69504) according to the manufacturer’s protocol for cultured cells. Library diversity was assessed by nested PCR amplification. The first PCR was performed using KAPA HiFi HotStart ReadyMix (ThermoFisher #NC0465187) with forward primer ctgagcaaagaccccaacgagaa and reverse primer gaaggcacaggtcgacaccagt under the following conditions: 98°C for 2 min; 10 cycles of 98°C for 10 sec, 55°C for 25 sec, 72°C for 30 sec; 72°C for 5 min. PCR products were purified using 1.5x SPRIselect beads (Beckman Coulter #A63880). Half of the eluate was used as template for a second PCR with forward primer ACACTCTTTCCCTACACGACGCTCTTCCGATCTNNNNNNGAGTAACCGTTGCTAGGAGAGA

CCATAT and reverse primer GTGACTGGAGTTCAGACGTGTGCTCTTCCGATCTcacaggtcgacaccagtctcatt under the following conditions: 98°C for 2 min, 10 cycles of 98°C for 10 sec, 58°C for 25 sec, 72°C for 30 sec, and 72°C for 5 min. Following 1.5x SPRI bead cleanup, the product was used as template for a third PCR to add Illumina sequencing adapters with indexed forward primer (P5) AATGATACGGCGACCACCGAGATCTACACgtacgattctACACTCTTTCCCTACACGACGCTCTT CCGATCT and indexed reverse primer (P7) CAAGCAGAAGACGGCATACGAGATttaccttgcaGTGACTGGAGTTCAGACGTGTGCTCTTCCG ATCT under the following conditions: 98°C for 2 min, 8 cycles of 98°C for 10 sec, 62°C for 20 sec, 72°C for 30 sec, and 72°C for 5 min. Libraries were purified using 1.5x SPRI beads and sequenced on an Illumina NextSeq 2000 with PE reads (89 cycles + 129 cycles) by the OHSU MPSSR. A total of 11,225,471 unique barcodes were identified in the library.

##### Lentivirus production and transduction

Lentivirus was produced by transfecting HEK293T LentiX cells (Clontech/Takara #632180) with pLARRY-EGFP-BCv1 transfer plasmid along with packaging plasmid psPAX2 and envelope plasmid pMD2.G using FuGENE 6 Transfection Reagent (Promega #E2691). psPAX2 (Addgene plasmid #12260) and pMD2.G were gifts from Didier Trono (Addgene plasmid #12259). HEK293T LentiX cells were cultured in DMEM (Gibco #11965-092) supplemented with 10% FBS and 1% penicillin-streptomycin (Gibco #15140122). Cells were seeded at 8e5 cells per well in 6-well plates and incubated overnight. Transfection complexes were prepared by combining Opti-MEM (Gibco #31985062) with FuGENE 6 per well, vortexing, and incubating for 5 minutes at room temperature. The transfer, packaging, and envelope plasmids were added to the FuGENE 6-Opti-MEM mixture, vortexed immediately, and incubated for 15-20 minutes at room temperature before adding dropwise to cells. Viral supernatant was harvested at 48 and 72 hours post-transfection, pooled, filtered through a 0.45 μm PES filter (Millipore Sigma #HPWP01300), and concentrated using Lenti-X Concentrator (Takara #631231) according to the manufacturer’s instructions. Briefly, Lenti-X Concentrator was combined with clarified supernatant at a 1:3 ratio and incubated at 4°C for 30 minutes. The mixture was centrifuged at 1,500 for 45 minutes at 4°C and the pellet was gently resuspended in complete PBS. Concentrated virus was aliquoted and stored at -80°C.

OCI-AML8227 cells were transduced with concentrated pLARRY lentivirus via spinoculation. Cells were seeded at 4e5 cells/mL in 24-well plates and supplemented with HEPES buffer (Gibco #15630-080) and polybrene (8 μg/mL; Millipore #TR-1003-G). Titrated volumes of concentrated lentivirus were added to achieve approximately 10% GFP infectivity. Plates were centrifuged at 800g for 90 minutes at 32°C with the brake disabled and then returned to the incubator overnight. The following day, cells were pelleted by centrifugation, resuspended in fresh culture medium, and returned to the incubator. On day 5 post-transduction, cells were split into two equal fractions. One fraction was treated with either DMSO or 0.5 μM venetoclax (MedChem Express #HY-15531) and incubated for 72 hours. The other fraction was immediately prepared for FACS. On day 8 post-transduction (72 hours post-treatment), the treated cell fractions were prepared for FACS. For FACS preparation, cells were centrifuged at 2,500 rpm for 3 minutes, washed with FACS buffer (PBS supplemented with 2% FBS), centrifuged again at 2,500 rpm for 3 minutes, resuspended in 100 μL FACS buffer, filtered through a 35 μm cell strainer cap, and sorted using a Sony SH800S cell sorter. GFP-positive cells were collected into 1.5 mL tubes on ice and resuspended to a final concentration of 1,000 cells/μL.

##### LARRY barcode amplification

LARRY barcodes and single-cell libraries were amplified using 10x Genomics single-cell cDNA libraries following an adapted protocol. Following Step 2.4 (cDNA QC & Quantification) of the Chromium Next GEM Single Cell 3’ Reagent Kits v3.1 protocol, amplified cDNA was used as template for barcode-specific PCR. The first PCR was performed using 2x KAPA HiFi HotStart ReadyMix (Roche #07958935001), cDNA template, 10x SI Primer (10x Genomics #PN-2000095), and 10 μM reverse primer TruseqR2_Larryv1. The reaction was cycled as follows: 95°C for 3 min, 98°C for 20 sec, 20-25 cycles of 98°C for 20 sec, 67°C for 15 sec, 72°C for 20 sec, and 72°C for 1 min. The optimal number of cycles was determined by monitoring exponential amplification via qPCR using an Applied Biosystems QuantStudio 6 (ThermoFisher). PCR products were purified using 1.8x MagBind TotalPure NGS beads (Omega Bio-tek #M1378-01), washed with fresh 80% ethanol, and eluted in nuclease-free water. The second PCR was performed using product from the first PCR along with 2x KAPA HiFi HotStart ReadyMix, template, 10x SI Primer, and indexed reverse primer from the Single Index Plate T Set A (10x Genomics #PN-1000213). The sample index well ID was recorded for each sample. The reaction was cycled as follows: 95°C for 3 min, 98°C for 20 sec, 10-15 cycles of 98°C for 20 sec, 54°C for 15 sec, 72°C for 20 sec, and 72°C for 1 min. PCR products were purified using 1.8x MagBind beads and eluted in nuclease-free water. Libraries were quantified using an Invitrogen Qubit Flex Fluorometer (Thermo Fisher Scientific #Q33326) and an Agilent TapeStation (Agilent #G2992A) and sequenced on an Illumina NextSeq Express (SP with 100 bp PE reads) by the OHSU MPSSR.

#### CRISPR

##### Sample preparation

CRISPR guide RNAs (gRNAs) in a pLentiCRISPR v2 backbone were obtained from GenScript. Lentivirus was produced by transfecting Lenti-X 293T cells (Clontech) with the SMARTvector transfer plasmid and packaging/pseudotyping plasmids. psPAX2 was a gift from Didier Trono (Addgene plasmid #12260). The supernatant containing lentivirus was collected after 48 hours of culture and filtered with a 0.45 μm filter. OCI-AML8227 cells were transduced with lentivirus via spinoculation in the presence of polybrene. Transduced cells were selected with 1 μg/mL puromycin to produce a stable cell line.

##### Validation

Cells were evaluated for cell surface marker expression after expansion for 2-4 weeks. Extra cells were set aside to extract cellular DNA using the DNeasy Blood and Tissue kit (Qiagen #69506) and then each gene of interest was amplified with PCR. Paramagnetic beads were used to purify the PCR DNA fragments (MagBio #AC-60001) and subsequently sequenced by Eurofins Genomics. Gene knockdown was validated using Tracking of Indels by Decomposition (TIDE)^18^. Inference of CRISPR Edits (ICE) was performed using the Synthego web tool (https://ice.synthego.com/#/). The PCR-amplification primers were designed in Geneious Prime and synthesized by Integrated DNA Technologies (Supplementary Table 1).

#### Drug sensitivity assays

##### Drug delivery

Drugs were delivered via printing using an HP 300e Digital Dispenser (HP #F0L56A). Each drug was plated on a 5-, 7-, or 8-point concentration curve starting with 0 μM, with exact drug concentrations dependent upon individual cell line sensitivity and the measurement assay in question.

##### MTS assay

Cells were cultured in 384-well plates with 50 μL media per well at an appropriate density to avoid over- or under-growth. A control row with only media was included. For MUTZ3, since some of the old media was used as part of the new media preparation, the media used for the control row was filtered to ensure there were no cells. To avoid edge effects, the outer wells were filled with DPBS. On day 3, cells were exposed to 5 μL (1:10 dilution) of CellTiter 96 AQueous One Solution Reagent (Promega #G358X) and incubated for 2-5 hours prior to reading with a BioTek Synergy 2 microplate reader (BioTek Instruments). At least 3 replicates were completed per cell line. Within each plate, each well was normalized to the average of the control row. Normalized matrices were exported and then averaged across replicates using Excel, generating a single matrix per cell line. Negative values were converted to zero and analyzed for drug synergy using the SynergyFinder web interface (https://synergyfinder.fimm.fi/) based on the Zero Interaction Potency (ZIP) reference model^19–21^.

##### Guava Nexin assay

Each patient sample tube was thawed at 37°C, diluted by dropwise adding 9 mL StemSpan SFEM-II (StemCell Technologies #09655), and allowed to rest for 10 minutes at room temperature before centrifugation and resuspension in DPBS. The Dead Cell Removal Kit (Miltenyi Biotec #130-090-101) was then used, typically with an MS column unless more than 10 million dead cells were detected. Healthy live cells were collected in the flow-through. Cells were suspended at 10,000-20,000 cells in 50 μL serum-free expansion media per well in a 96-well plate (StemSpan SFEM-II [StemCell Technologies #09605], 1:20 StemSpan CD34+ Expansion Supplement [StemCell Technologies #02691], 1:500 UM729 [StemCell Technologies #72332]). Using an HP 300e Digital Dispenser (HP #F0L56A), 5 concentrations (including 0 μM) of venetoclax and/or DB2313 were printed in a 5×5 matrix with 3 replicates per condition and incubated for 72 hours. On day 3, Guava Nexin Reagent (Cytek #4500-0455) was applied at a 1:1 ratio for 15-30 minutes at room temperature. The Guava easyCyte HT System was used to collect 2,000 events per well, with 3 seconds of paddle mixing per well. Three replicates per patient were collected, although a clog occurred with two plates, resulting in two patients having only 1 replicate with accurate readings. Gating of cell populations was performed using guavaSoft 3.4, with viability measured as cells that were negative for both Annexin V-PE and 7-AAD. Final analyses and plots were performed using the SynergyFinder web interface (https://synergyfinder.fimm.fi/) based on the ZIP reference model^19–21^. One sample was excluded from downstream analyses due to widespread cell death (18-00105). Viability data was averaged across all replicates and normalized to the untreated well to obtain final 5×5 matrices of viability values per patient, which were then analyzed for drug synergy. Prior to calculating average synergy across all patients, all replicates within each patient were averaged so that patients with more replicates would not be overrepresented.

### Sequencing

#### Genetrails Hematologic Malignancies Sequencing Panel

Genetic profiling was completed by the Knight Diagnostics Laboratory using standard clinical workflows.

#### Single-cell RNA sequencing

For patient samples, frozen viable cells stored in liquid nitrogen were thawed at 37°C before adding dropwise thaw media (49.5% FBS, 49.5% X-VIVO 10 [Lonza #04-380Q]) and allowed to rest for 10 minutes at room temperature before centrifugation and resuspension in DPBS. The Dead Cell Removal Kit (Miltenyi Biotec #130-090-101) was then used, typically with an MS column unless more than 10 million dead cells were detected. Healthy live cells were collected in the flow-through and fixed using the Scale FixKit v1, then stored at -80°C. Fixed cells were thawed at room temperature and multiplexed (12 or 16 samples) to generate scRNA-seq libraries using the ScaleBio 3-level scRNA kit (v1 or v2). For the primary OCI-AML8227 libraries, cells were cultured with DMSO or 1 μM venetoclax for 72 hours prior to collection and scRNA library generation using the Chromium Single Cell RNA Library and Dual Index Next GEM Single Cell 3’ Reagent Kits v3.1 (10x Genomics #PN-1000268), and the clonal barcoding libraries were prepared following the LARRY protocol as described above^17^. Libraries were sequenced with 100 bp PE sequencing by the OHSU MPSSR.

#### Single-cell ATAC-seq

OCI-AML8227 cells were treated with DMSO or 1 μM venetoclax for 24 hours before assessment of chromatin accessibility. Single-cell ATAC-seq was performed using the s3ATAC-seq protocol as previously described^22^. Briefly, nuclei were isolated from OCI-AML8227 cells using nuclei isolation buffer (10 mM HEPES-KOH pH 7.2 [Fisher Scientific #BP310-500 and Sigma Aldrich #1050121000], 10 mM NaCl [Fisher Scientific #S271-3], 3 mM MgCl₂ [Fisher Scientific #AC223210010], 0.1% IGEPAL-CA630 [Sigma Aldrich #I3021], 0.1% Tween-20 [Sigma Aldrich #P-7949], and EDTA-free Pierce protease inhibitor [ThermoFisher #A32955]). Nuclei were tagmented in tagmentation buffer (132 mM TAPS pH 8.5 [Sigma Aldrich #T0647], 264 mM potassium acetate [Sigma Aldrich #P1190], 40 mM magnesium acetate tetrahydrate [Sigma Aldrich #M5661], 12 mM D-Glucosamine sulfate [Fisher Scientific #AAJ6627106], and N,N-Dimethylformamide [Sigma Aldrich #D4551]) using Tn5 transposase (Scale Biosciences). During tagmentation, nuclei were incubated for 30 minutes at 42°C while shaking at 300 RPM on an Eppendorf ThermoMixer C. Tagmented nuclei were pooled, stained with DAPI (Cell Signaling Technologies #4083), and sorted into a 96-well plate using a Sony SH800S cell sorter. Nuclear proteins were denatured by incubation at 55°C for 15 minutes on an Eppendorf Mastercycler nexus with the lid heated to 90°C. Linear extension and adapter switching were performed in nuclei resuspended in 300 nM LNA oligo, 1.5% Triton-X100 (Sigma Aldrich #T8787), and VeraSeq PCR Mix (Qiagen #P7610L) on an Eppendorf Mastercycler nexus. qPCR was performed using VeraSeq Buffer II (Qiagen #B7102), 10 mM dNTPs (ThermoFisher #R0192), and VeraSeq Ultra Polymerase (Qiagen #P7520L). Libraries were cleaned using the DNA Clean & Concentrator-5 kit (Zymo Research #D4013) and then were size-selected using MagBind TotalPure NGS Binding Beads (Omega Biotech #M1378-01). Cleaned libraries were quantified using an Invitrogen Qubit and an Agilent TapeStation. Libraries were diluted to 600 pM and then sequenced with 100 bp PE on an Illumina NextSeq 2000.

### Analysis

#### Beat AML cohort

Patient data from the BeatAML 2.0 cohort dataset was obtained according to IRB 4422 biorepository study^8,23^. For analysis of cohort-level data, samples collected from patients who were in remission, relapse, or had a history of myelodysplastic syndrome at the time of collection were excluded, resulting in 645 AML samples analyzed. When enriching for samples above the 33% threshold, the total number of samples analyzed was n = 289. For mutation calls, variants were only called as a relevant mutation if they were "possibly_damaging" or "probably_damaging" by PolyPhen, or "deleterious" or "deleterious_low_confidence" by SIFT. Gene mutations shown were detected in at least five samples. Markers were binarized (present/absent) and those shown were detected in at least 10 samples. Drugs shown were measured in at least 10 samples per group and significance was determined by multiple t-tests adjusted with Benjamini-Hochberg correction.

#### Flow cytometry

After staining, cells were washed with stain buffer and analyzed using a Symphony A5 instrument. Flow cytometry data was analyzed using FlowJo (version 10). Live single cells were isolated using Live/Dead Blue, forward scatter, and side scatter. Gating of individual markers was guided by unstained, single stain, and fluorescence-minus-one (FMO) control stains. Venetoclax showed autofluorescence in the APC channel and DB2313 showed autofluorescence in the UV channel. For DB2313 experiments, forward and side scatter alone were used for discrimination of live single cells. Cell counts per subpopulation were calculated by multiplying live cell counts from the TC20 by the proportion of each subpopulation obtained from gating in FlowJo.

#### Phosphoproteomics

##### TMT Global Proteomics Data Processing

All Thermo RAW files were processed using mzRefinery to correct for mass calibration errors, and then spectra were searched with MS-GF+ v9881 to match against the Uniprot reference protein sequence database from April 2018 (71,599 entries)^24–26^. A partially tryptic search was used with a ± 20 ppm parent ion mass tolerance. A reversed sequence decoy database approach was used for false discovery rate calculation. MS-GF+ considered static carbamidomethylation (+57.0215 Da) on Cys residues and TMT modification (+229.1629 Da) on the peptide N terminus and Lys residues, and dynamic oxidation (+15.9949 Da) on Met residues. The resulting peptide identifications were filtered to a 1% false discovery rate at the unique peptide level. A sequence coverage minimum of 6 per 1000 amino acids was used to maintain a 1% FDR at the protein level after rollup by parsimonious inference. The intensities of TMT 16 reporter ions were extracted using MASIC software^27^. Extracted intensities were then linked to PSMs passing the confidence thresholds described above. Relative protein abundance was calculated as the ratio of sample abundance to reference channel abundance, using the summed reporter ion intensities from peptides that could be uniquely mapped to a gene. The relative abundances were log2 transformed and zero-centered for each gene to obtain final relative abundance values.

##### TMT Phosphoproteomics Data Processing

IMAC enriched fraction datasets were searched as described above with the addition of a dynamic phosphorylation (+79.9663 Da) modification on Ser, Thr, or Tyr residues. The phosphoproteomic data were further processed with the Ascore algorithm for phosphorylation site localization, and the top-scoring assignments were reported^28^. The relative abundances were log2 transformed and zero-centered for each phosphosite to get final relative abundance values.

##### Causal network analysis

Normalized global and phosphoproteomics datasets underwent causal network analysis using CausalPath^29^. Two conditions were compared using fold change of mean values. The threshold for data significance was set to 2 for changes in protein levels in the global proteomics analysis and 1.75 for changes in both global protein levels and phosphoprotein levels in the phosphoproteomics analysis. Relationships meeting these thresholds were visualized using the CausalPath web interface (https://causalpath.cs.umb.edu/).

#### Single-cell RNA seq

##### Pre-processing

All libraries were aligned to the hg38 human genome. For ScaleBio libraries, the ScaleBio scRNA pipeline (https://github.com/ScaleBio/ScaleRna) was used for demultiplexing BCL files into FASTQ files. For OCI-AML8227 clonal barcoding libraries, a significantly higher number of cells were sampled in the DMSO 72-hour sample with less coverage per cell on average. To equalize the opportunity to identify dual-timepoint clones between DMSO and venetoclax treatment conditions, DMSO-treated samples were downsampled prior to genome alignment at the FASTQ level. These clonal barcoding libraries were then processed using cloneRanger, a pipeline that incorporates 10x Genomics Cell Ranger (v7.1.0) for read alignment and feature quantification, followed by clonal barcode processing and cell assignment (https://github.com/dfernandezperez/cloneRanger)^30^. LARRY clonal barcodes were processed using the following parameters in cloneRanger: Hamming distance of 3, minimum read count per molecule (reads_cutoff) of 10, and minimum UMI count (umi_cutoff) of 3. Clonal barcode patterns were specified according to LARRY v1 barcodes used for library construction (Addgene LARRYv1 library #140024), which consist of a 40-nucleotide sequence containing 28 random nucleotides evenly interspersed with six pairs of fixed nucleotides^17^.

##### Quality filtering

The Seurat package in R was used to import all samples as individual Seurat objects. Genes appearing in fewer than 3 cells per sample were removed, and >500 genes were required to retain a cell. For 10x Genomics libraries, cells with >10% mitochondrial genes were excluded^31^. For patient samples, additional thresholds of <2% mitochondrial genes and 1,000–25,000 transcripts were applied. Doublets were removed if called by both scDblFinder (https://github.com/plger/scDblFinder)^32^ and the hybrid score from the scds package (https://github.com/kostkalab/scds)^33^. For OCI-AML8227 libraries, cells deviating (z-score >3) from the regression line of transcript by gene counts across all samples in that set were additionally removed. Cells considered outliers by either gene counts or transcript counts (z-score >3) were also excluded. Doublets identified by clonal barcodes were removed. Singlet cells contained ≥10-fold more reads of the primary barcode than any secondary barcode, while doublet cells showed <10-fold difference between primary and secondary barcodes. Doublet pairs appearing at both timepoints within a treatment group but absent as primary singlets were retained, interpreting these as rare double-insertion events.

##### Mapping and annotation

For each experiment, all libraries were merged into a single object and mapped into the UMAP space of a healthy single-cell transcriptomic atlas of hematopoiesis^34^. The BoneMarrowMap package in R (https://github.com/andygxzeng/BoneMarrowMap) was used to predict cell type and pseudotime for each cell, discarding cells that did not receive a cell type assignment.

##### Analysis and plots

For patient samples, cell proportions were calculated using all mapped cells obtained for each sample. In UMAPs, to prevent overrepresentation of samples with higher cell counts, an equal number of cells from each sample (∼450 cells) was used. For OCI-AML8227 cells, cell proportions were calculated using all clonally barcoded cells (∼80% of each library). The analysis was downsampled to equalize the number of paired clones per treatment condition, and significance was tested using a Kolmogorov-Smirnov test. Since clonal relationships could be 1:1, 1:many, many:1, or many:many, each unique cell pair was recorded. Consequently, some individual cells appear in multiple pairs. To equalize the opportunity to identify clones between control and treated samples, the analysis was downsampled within each timepoint to the same number of clonally barcoded cells (∼3,000 cells per 0-hour library, ∼4,100 cells per 72-hour library).

For OCI-AML8227 samples, the Python implementation of SCENIC (pySCENIC) was run on normalized counts from Seurat-processed cells that passed quality filtering^35,36^. Gene regulatory network (GRN) inference was performed with the GRN module using the default GRNBoost2 algorithm and a transcription factor list provided in the SCENIC documentation. Predicted regulons were identified with the cisTarget module using motif annotations and genome ranking databases also provided in the SCENIC documentation. Predicted regulon activity was quantified for each cell using the AUCell module. To compare regulon activity between DMSO and venetoclax treatment conditions at each timepoint, Kolmogorov-Smirnov tests were performed on AUCell score distributions, and p-values were adjusted using Benjamini-Hochberg correction. AUCell scores were plotted along pseudotime for each condition and timepoint, and smoothed trend lines were used to summarize predicted transcription factor activity and infer the cell types and pseudotime intervals where activity diverged.

#### Single-cell ATAC-seq

##### Pre-processing

Libraries were demultiplexed with Tn5, i5, and i7 barcodes and the Tn5 tagmentation plate position using a custom script. Demultiplexed FASTQ files were aligned to the human genome (hg38) using BWA-MEM^37,38^. Aligned reads were filtered to exclude unmapped or improperly paired reads, as well as reads aligned to mitochondrial DNA, chromosome Y, or noncanonical contigs using SAMtools^39^. Fragments between 20–10,000 bp were retained. Duplicates were removed using a custom script.

##### Analysis

ArchR was used for cell quality filtering, normalization, dimensionality reduction, peak calling, unsupervised clustering, gene accessibility analysis, differential accessibility analysis, and motif enrichment^40^. Cells passing quality thresholds (transcription start site enrichment ≥5, unique fragments ≥1,500) were retained for further analysis. The ArchR implementation of MACS2 was used for calling peaks^41^. Cells were also clustered based on a peak matrix derived from the reproducible peak set generated by ArchR. To infer cell types present in these clusters, gene accessibility across cells was assessed using ArchR gene scores for known marker genes, including CD34 and CD38 for hematopoietic stem and progenitor cells, MPO and LYZ for myeloid progenitors, and CD14 for monocytes. For differential accessibility analysis at each condition and timepoint, cells in mature monocytic clusters were compared to cells in more primitive progenitor-like clusters. Motif enrichment was assessed separately in significantly upregulated and downregulated differential peaks (FDR ≤0.1, absolute log2 fold change ≥0.25), and motifs were ranked according to enrichment significance.

#### CRISPR

After calculating cell counts as described in the flow cytometry section above, total cell counts for each knockout cell line were normalized to the DMSO average (set to 100). Statistical significance was calculated in Prism using two-way ANOVA with Holm-Šidák correction. When data failed at least two of four normality tests (D’Agostino-Pearson omnibus, Anderson-Darling, Shapiro-Wilk, and Kolmogorov-Smirnov), values were transformed to fit residuals to a Gaussian distribution prior to ANOVA.

### Quantification and Statistical Analysis

Values are represented as the mean, and error bars are the SEM unless otherwise stated. Prism (version 10.6.1; Prism Software Corp.) or R was used for statistical analyses. Significance was tested using Student’s t-test or ordinary one-way ANOVA followed by Holm-Šidák post-test correction unless otherwise stated.

### Data Availability

All raw and processed sequencing data is available upon request.

## RESULTS

### Stem–monocytic AML represents a transcriptionally distinct subtype with reduced sensitivity to BCL2 inhibition

The therapeutic impact of BCL2 inhibition on patients harboring both immature and differentiated leukemic characteristics remains poorly characterized. To address this, we analyzed molecular data from the Beat AML cohort, one of the largest primary patient AML repositories containing paired *ex vivo* drug sensitivity, RNA sequencing (RNA-seq), whole exome, and targeted DNA sequencing (DNA-seq)^8,23^. We scored each patient sample using previously described cell type gene expression signatures to determine enrichment for transcriptional differentiation states (Figure 1a; Supplementary Figure 1a). Consistent with prior reports, patient samples demonstrated disease characteristics across the spectrum of myeloid differentiation^5,6,8^. However, a substantial population (26%) showed enrichment for both HSC- and monocytic-like transcriptional signatures, which we termed stem-monocytic AMLs (Figure 1b). Given the established sensitivity of immature AML to BCL2 inhibition with venetoclax, we analyzed paired *ex vivo* drug-sensitivity data from these patient samples^9–11^. Consistent with prior reports, patient samples enriched with HSC-like transcriptional signatures were exquisitely sensitive to venetoclax as compared to those enriched with monocyte-like signatures (Figure 1c; Supplementary Figure 1b). Critically, however, the patient population enriched with both HSC-and monocyte-like transcriptional features was minimally responsive to venetoclax, comparable to populations with predominantly monocyte-like disease. We extended this analysis to additional therapeutic agents and found that the *ex vivo* drug-response profile of patients with combined HSC- and monocyte-like features closely resembled that of monocyte-like disease (R² = 0.975; Supplementary Figure 1c-e). Together, these findings indicate that stem-monocytic AML exhibits functional characteristics that resemble monocyte-like AML, particularly with respect to BCL2 inhibitor sensitivity.

**Figure 1:**
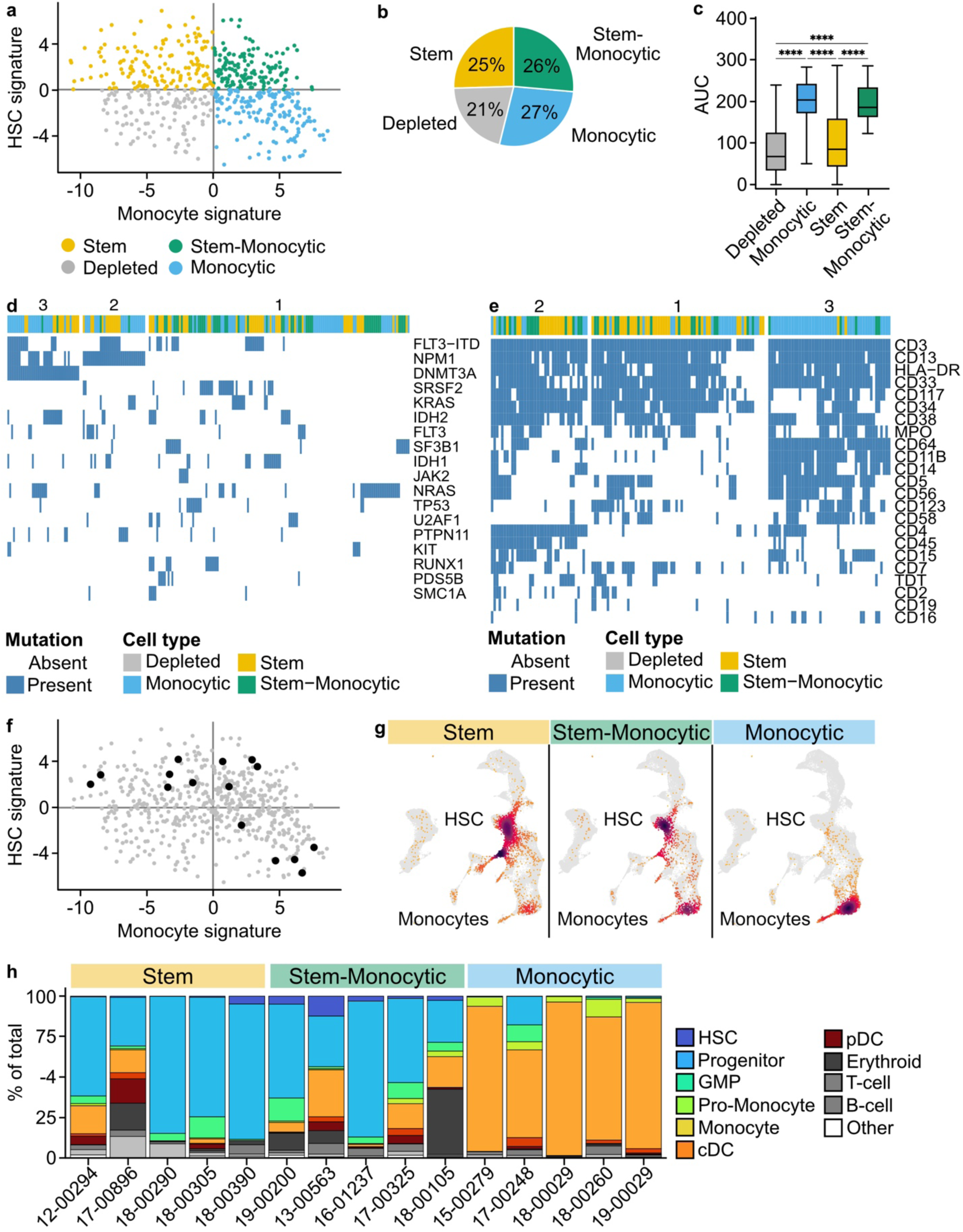
Stem-monocytic AML is a functionally distinct subtype with reduced sensitivity to BCL2 inhibition. **a.** Cell type transcriptional signatures were calculated using previously described methods for 652 patients in the BeatAML cohort^8^. The first principal component of the hematopoietic stem cell (HSC) and monocyte signatures is visualized, where each point represents a single patient. Patients were stratified based on HSC and monocyte transcriptional signatures. The four colors correspond to the transcriptional subtypes: stem, depleted, monocytic, and stem-monocytic. **b.** Pie chart displaying the proportions of patients classified in each transcriptional subtype. **c.** Primary AML blasts were cultured for 72 hours along a 7-point dose curve with venetoclax. Cell viability was assessed by CellTiter Aqueous colorimetric assay. Area under the curve (AUC) values from the dose-response curves are shown for each transcriptional subtype. Patients within the top 33% of each transcriptional subtype were selected for analysis. Significance was evaluated using Brown-Forsythe and Welch ANOVA tests followed by Dunnett T3 post-test correction to account for non-normal distribution of the data. **d.** Mutational profiles for each patient as determined by targeted genetic sequencing. Patients within the top 33% of each transcriptional subtype were selected for analysis. Each patient is color-coded according to subtype. **e.** Immunophenotypic profiles of patients within the top 33% of each transcriptional subtype. **f.** Cell type transcriptional signatures of the 15 primary AML samples chosen for scRNA-seq. **g.** UMAP dimensional reduction and projection of single-cell gene expression profiles. Single-cell density is shown for each transcriptional subtype. UMAP space and cell assignment were determined by mapping transcriptional profiles to a single-cell reference atlas^34^. **h.** Stacked bar plots displaying the proportion of lineages within each sample in panel g. ns = not significant; * = p < 0.05; ** = p < 0.01; *** = p < 0.001; **** = p < 0.0001.

Conventional clinical techniques, including genotyping and immunophenotyping, were evaluated for their utility in stratifying these populations. K-means clustering failed to identify consistent patterns among the stem-monocytic population, which more closely resembled the immunophenotypic profile of stem AML (Figure 1d,e; Supplementary Figure 2). These findings indicate that conventional diagnostic approaches may be insufficient for identifying these patients. To further characterize the transcriptional landscape of these patient populations, we performed single-cell RNA seq on 15 patient samples from this cohort (Figure 1f). We mapped transcriptional profiles to a single-cell reference atlas to identify cell types^34^. The highest density of leukemic cells in primarily HSC-like and monocyte-like samples aligned with their respective transcriptional signatures in the reference atlas (Figure 1g,h). In contrast, patient samples enriched for both transcriptional signatures were broadly distributed across differentiation states rather than localized to discrete HSC- or monocyte-like populations. These observations establish stem-monocytic AML as a transcriptionally distinct entity characterized by heterogeneous leukemic cell populations that exhibit resistance to BCL2 inhibition comparable to monocytic disease despite immunophenotypic similarity to immature AML.

### The OCI-AML8227 cell line recapitulates patient responses to venetoclax response and models stem-monocytic AML

In order to characterize the functional consequences of differentiation heterogeneity observed in stem-monocytic AMLs, we sought a tractable model system with similar characteristics. Functional studies in primary AML samples are technically challenging, and most immortalized human AML cell lines exhibit homogeneous differentiation states. Therefore, we used the OCI-AML8227 cell line, a patient-derived AML culture system that maintains both clonogenic progenitors and terminally differentiated cells^13^. To identify differentiation states in OCI-AML8227, we developed a panel of four myeloid surface markers that together reveal six primary subpopulations reflecting normal myeloid development (Figure 2a; Supplementary Figure 3a). Notably, immature leukemic cells express CD34, whereas differentiated populations express monocyte-associated markers CD64 and CD14. Targeted DNA-seq revealed pathogenic mutations in *ASXL1*, *NRAS*, and *SRSF2*, consistent with AML arising from myelodysplastic syndrome or chronic myelomonocytic leukemia (Supplementary Table 2). These findings support the use of OCI-AML8227 as an appropriate model for stem-monocytic AML.

**Figure 2:**
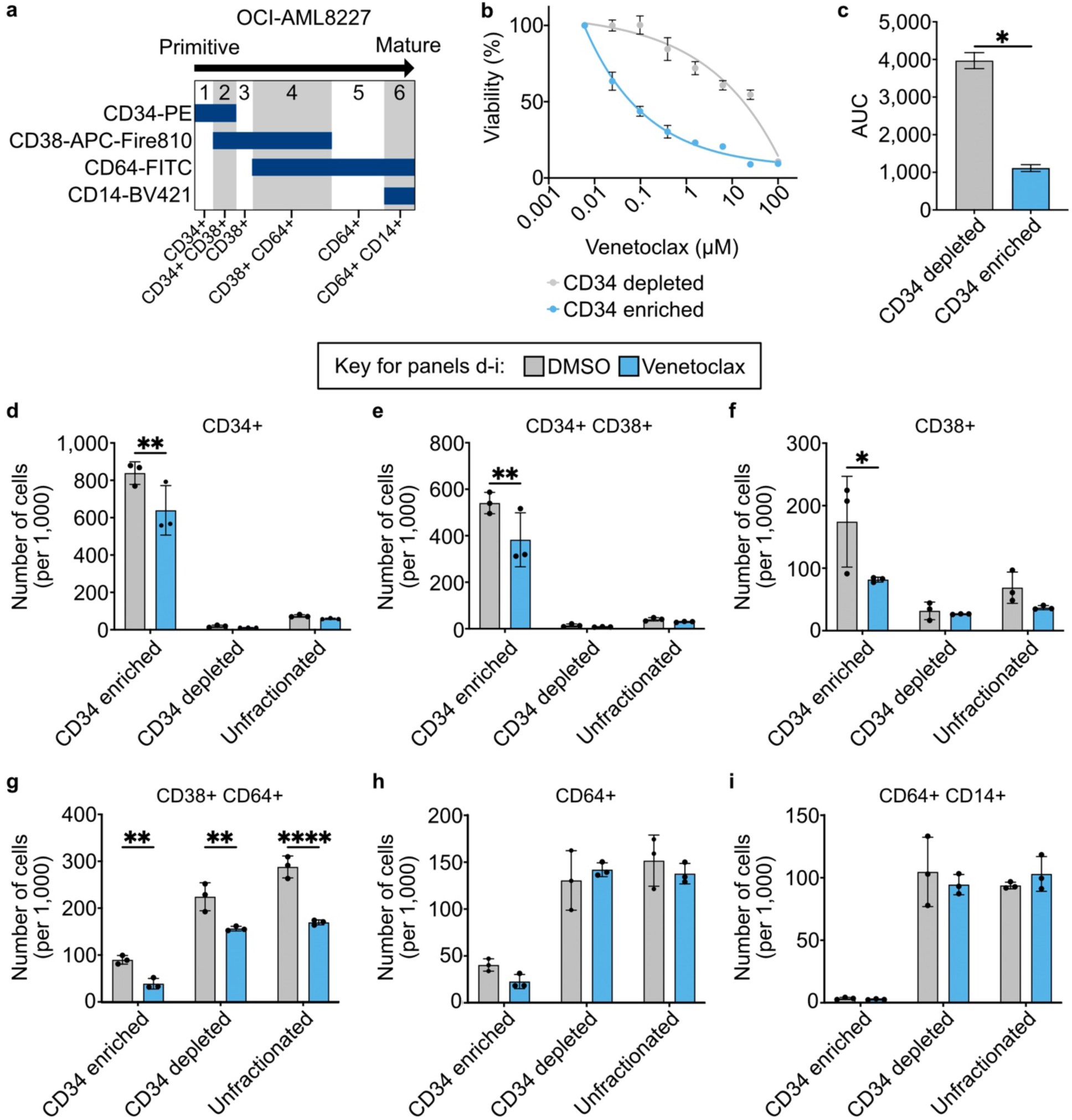
The OCI-AML8227 cell line model serves as a model of stem-monocytic AML. **a.** Schematic of differentiation heterogeneity in the OCI-AML8227 cell line model. This cell line produces leukemic blasts expressing immature myeloid markers (CD34) as well as differentiated monocytic markers (CD64 and CD14). CD38 is a transient myeloid differentiation marker, which is initially expressed in early myeloid progenitor cells. **b.** OCI-AML8227 cells were immunomagnetically fractionated by CD34 expression and cultured in triplicate for 96 hours along an 8-point dose curve with venetoclax. Cell viability was assessed by CellTiter Aqueous colorimetric assay. **c.** AUC values from the dose-response curves for each immunophenotypic population. Significance was evaluated using a two-tailed t-test. **d–i.** Cell surface expression of CD34, CD38, CD64, and CD14 in OCI-AML8227 cells cultured in triplicate for 72 hours with 1 μM venetoclax or an equivalent volume of DMSO. Cells were analyzed either as immunomagnetically fractionated populations (CD34-enriched and CD34-depleted) or as unfractionated cells. Significance was evaluated using ordinary one-way ANOVA followed by Holm-Šidák post-test correction. ns = not significant; * = p < 0.05; ** = p < 0.01; *** = p < 0.001; **** = p < 0.0001.

To assess the impact of differentiation state on venetoclax response, we immunomagnetically fractionated leukemic cells based on CD34 surface expression. Consistent with prior studies, CD34-enriched OCI-AML8227 cells were highly sensitive to venetoclax compared to CD34-depleted populations (Figure 2b,c). To investigate the mechanisms underlying this differential response, we performed phosphoproteomics following venetoclax treatment in these CD34-fractionated populations (Supplementary Figure 4; Supplementary Tables 3-5). Network-based analysis of global proteomic profiling revealed disruption of transcription factor regulators governing apoptosis and the cell cycle following BCL2 inhibition^29,42^. Specifically, CD34-enriched leukemic cells exhibited upregulation of a SP1-centered network and depletion of a MYC-centered network relative to CD34-depleted populations. Analysis additionally revealed modest but significant enrichment of BCL3, a transcriptional target regulated by both PU.1 and MYC. Analysis of the phosphoproteomic profiling data further demonstrated disruption of cell cycle regulators, including depletion of a CDK2-T160t-centered network and concurrent upregulation of a MAPK3-Y204y-centered network. These findings further support OCI-AML8227 as a model system for venetoclax response studies in this patient population and suggest that immature leukemic cells exhibit sensitivity to BCL2 inhibition mediated by disruption of central cell cycle and apoptotic regulators.

To understand the differentiation dynamics in response to BCL2 inhibition, we performed immunophenotypic profiling following venetoclax treatment. We compared immunomagnetically fractionated populations (CD34-enriched and CD34-depleted) with unfractionated cells (Figure 2d-i). Relative to DMSO-treated controls, venetoclax significantly depleted the four most immature subpopulations while sparing the two most differentiated subpopulations, consistent with venetoclax resistance observed in monocytic patient samples. The similar responses of unfractionated and CD34-depleted populations suggest that differentiation dynamics in stem-monocytic AML mirror those of monocyte-like disease.

### Lineage tracing demonstrates preferential survival of monocytic populations following venetoclax treatment

To define how clonal differentiation state influences sensitivity to BCL2 inhibition, we performed single-cell barcoding to track individual AML clones during venetoclax treatment^17^ (Figure 3a). Two OCI-AML8227 populations were individually transduced with a lentiviral barcode library, assigning unique identifiers to individual cells. Following a five-day recovery period to allow cellular recovery and outgrowth, each barcoded population was treated with either DMSO or venetoclax and split into two subpopulations: one was immediately processed for scRNA-seq (0-hour time point), and the other was cultured for an additional 72 hours before sequencing. Within each drug condition, unique barcodes detected at both time points (paired barcodes) were used to reconstruct clonal trajectories and identify populations that persisted or were depleted during treatment. This approach enabled direct linkage of transcriptional state with clonal survival and differentiation dynamics in response to BCL2 blockade.

**Figure 3:**
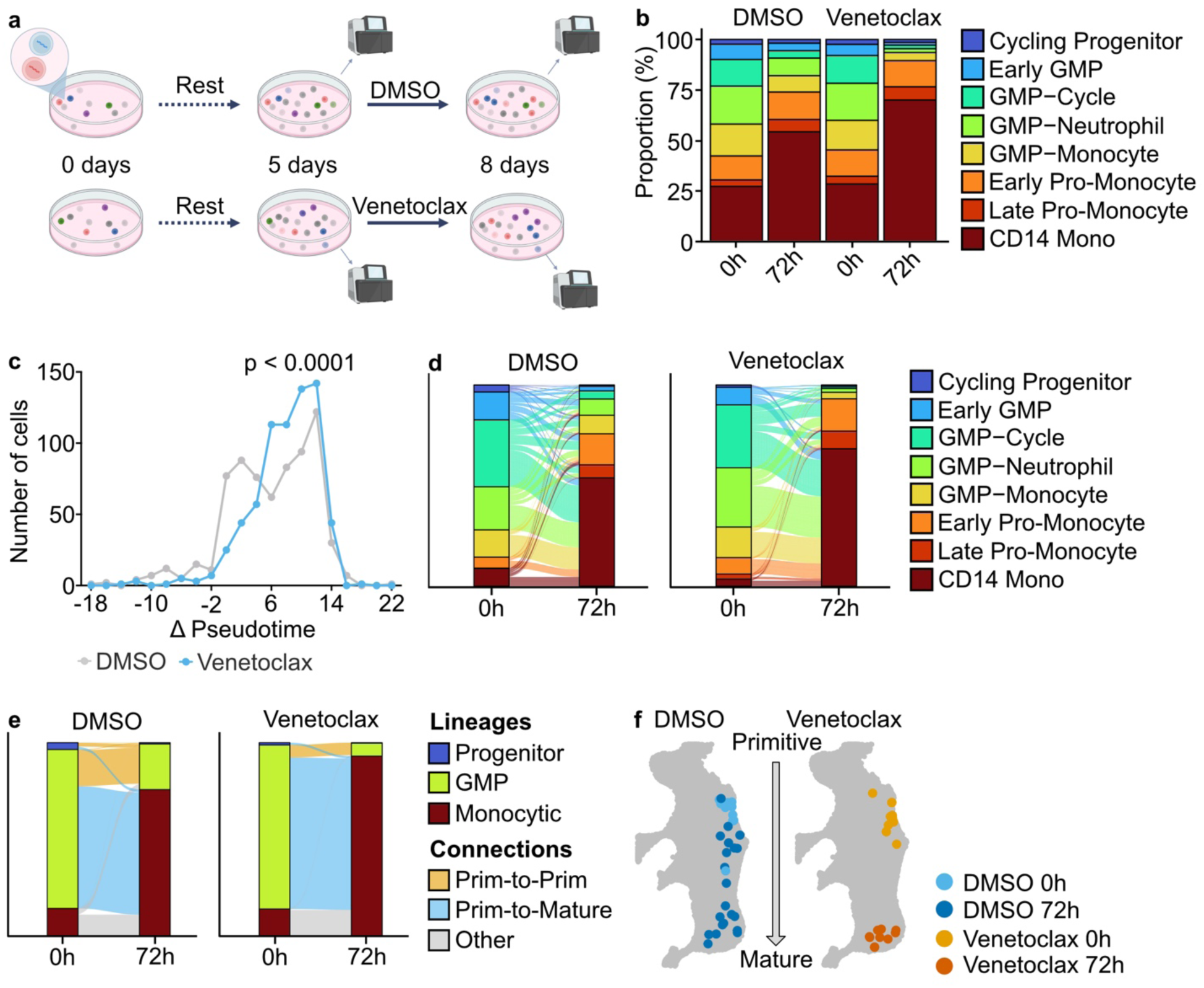
Lineage tracing demonstrates preferential survival of monocytic populations following venetoclax treatment. **a.** Schematic of lineage tracing experiments. Two OCI-AML8227 populations were individually transduced with a lentiviral barcode library by spinoculation. Following a five-day recovery period to allow cellular recovery and outgrowth, each barcoded population was treated with DMSO or 1 μM venetoclax and split into two subpopulations: one was immediately processed for scRNA-seq (0-hour time point) and the other was cultured for 72 hours before sequencing. Image was created with BioRender. **b.** Cell type assignments determined by mapping transcriptional profiles to a single-cell reference atlas^34^. Stacked bar plots display the proportion of cell types within each sample. **c.** Pseudotime analysis was performed to understand each cell’s relative position along the myeloid differentiation trajectory. The change in pseudotime (Δ pseudotime) was calculated for each paired barcode between the initial and 72-hour time points. Significance was evaluated using a Kolmogorov–Smirnov test. **d, e.** Sankey plots depicting cell (d) type and (e) lineage transitions of paired barcodes from the initial to the 72-hour time point. **f.** UMAP dimensional reduction and projection of single-cell gene expression profiles 0 hours or 72 hours following drug treatment with DMSO or 1 μM venetoclax. Gray dots represent the reference hematopoietic cells whereas the colored dots represent the drug-treated cells. ns = not significant; * = p < 0.05; ** = p < 0.01; *** = p < 0.001; **** = p < 0.0001.

We next examined how clonal composition and differentiation state changed following venetoclax exposure. Untreated OCI-AML8227 cells consisted largely of eight cell types, ranging from immature cycling progenitor- and GMP-like cells to mature pro-monocytic- and monocytic-like cells (Supplementary Figure 3b). Cell-type composition between treatment groups was similar at 0 hours. However, at 72 hours, venetoclax-treated cells showed an increased proportion of monocytic cells and a concurrent reduction of GMP cells (Figure 3b). To understand the clonal dynamics driving this difference, we identified unique clonal barcodes present at both time points within each treatment condition, representing ancestor-descendant relationships. To quantify differentiation shifts over time, we measured the change in pseudotime between 0 and 72 hours (Δpseudotime) for each clonally linked pair of cells (Figure 3c). Venetoclax treatment caused a shift in Δpseudotime values consistent with increased maturation. To determine which cells contributed to this increase in pseudotime, we examined direct connections across both discrete cell types and lineage groups (Figure 3d,e). Venetoclax treatment reduced the number of immature-to-immature relationships and concurrently increased the number of immature-to-mature relationships. This effect was particularly pronounced in clones originating from the most immature cell type, cycling progenitors. Whereas DMSO-treated cycling progenitor-like cells were associated with both immature and mature descendants, cells exposed to venetoclax were exclusively associated with mature descendants (Figure 3f). Together, these analyses demonstrate that venetoclax selectively eliminates self-renewing immature clones while selecting for expansion of differentiating clones. Furthermore, because OCI-AML8227 maintains a hematopoietic-like hierarchy and recapitulates validated responses to venetoclax, these results support its use as an appropriate model for stem-monocytic AML.

### Venetoclax-resistant blast populations are characterized by increased activity of myeloid differentiation regulators

To identify molecular regulators underlying the persistence of monocytic leukemic cells following venetoclax treatment, we analyzed predicted transcription factor activity from single-cell gene expression profiles generated in lineage tracing experiments. Comparisons of activity distributions between drug treatment conditions at both the initial and 72-hour time points identified 62 transcription factors with higher predicted activity in venetoclax-treated cells, including BACH1, CEBP family members, and PU.1 (encoded by *SPI1*; Figure 4a,b; Supplementary Figure 5a; Supplementary Table 5)^36^. To assess how activity of these transcription factors varied with differentiation, we examined predicted activity across pseudotime, which reflects each cell’s position along the myeloid differentiation continuum (Figure 4c; Supplementary Figure 5b,c). The activity of these factors increased with advancing differentiation across all treatment conditions, reflecting their role in myeloid maturation. Notably, venetoclax treatment at 72 hours was associated with sustained high activity of these transcription factors, suggesting that their maintenance may contribute to the emergence of monocytic, venetoclax-resistant AML populations (Figure 4b; Supplementary Figure 5). Together, these results indicate that stem-monocytic AML cells with limited venetoclax sensitivity are characterized by enrichment of transcription factors governing myeloid differentiation programs.

**Figure 4:**
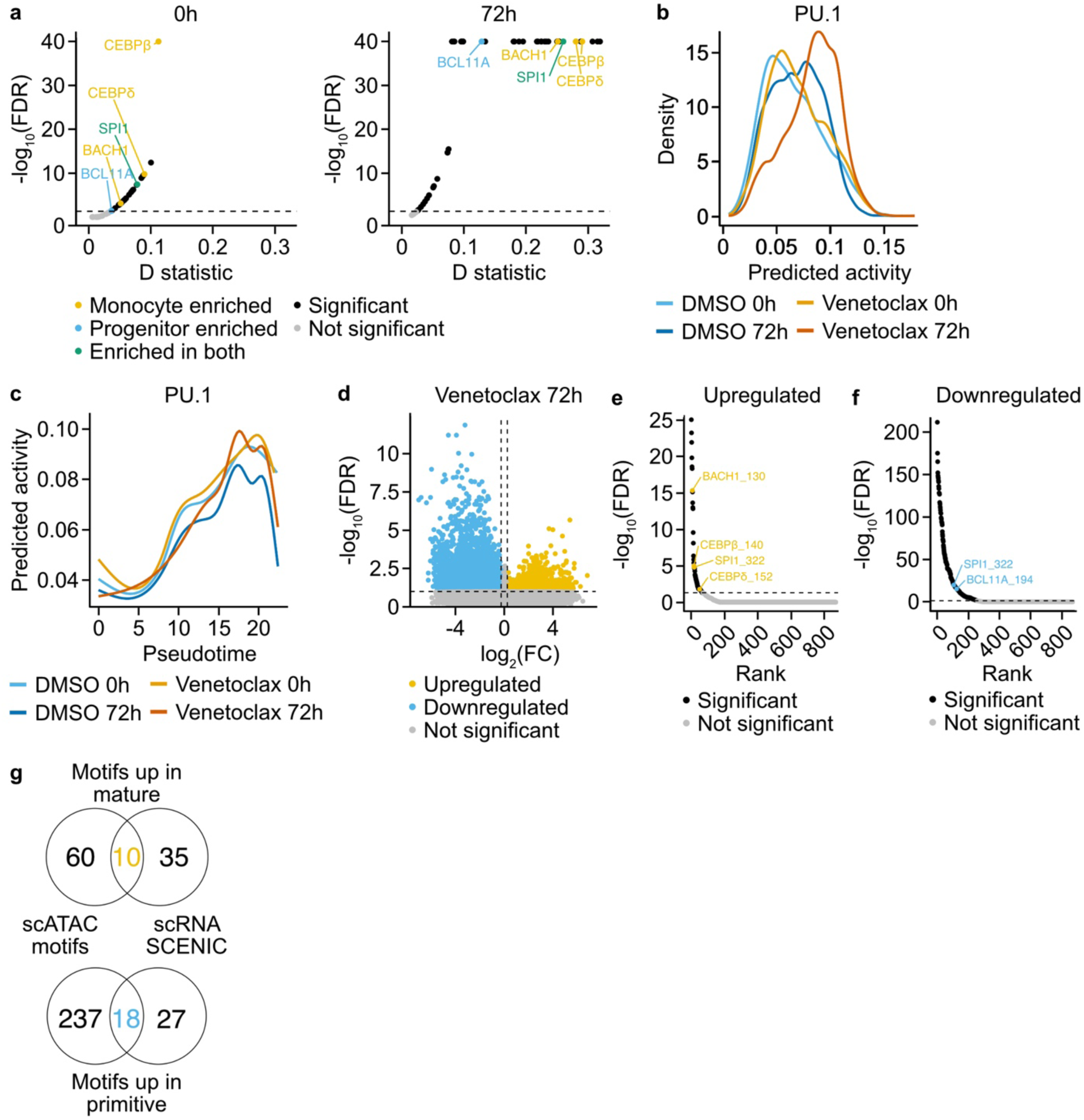
Venetoclax-resistant blast populations are characterized by increased activity of myeloid differentiation regulators. **a.** Predicted transcription factor activity was evaluated in single-cell RNA-seq data presented in Figure 3. Significantly enriched transcription factors were identified and denoted as enriched in progenitor (blue), monocytic (gold), or both (green) populations. Gray dots represent transcription factors that did not reach statistical significance (FDR < 0.05). **b.** Predicted activity of PU.1 visualized by each time point and drug condition. **c.** Pseudotime values generated in Figure 3 were used to determine each cell’s relative position along the myeloid differentiation trajectory. Predicted activity of PU.1 is visualized along its pseudotime trajectory in each drug condition. **d.** OCI-AML8227 cells were cultured with DMSO or 1 μM venetoclax and processed at either 0 or 72 hours for single-cell ATAC-seq. Differential peaks were identified from open chromatin regions in each condition. Significant peaks were compared between monocytic and progenitor clusters. Gray dots represent motifs that did not reach statistical significance (FDR < 0.05). **e, f.** Transcription factor motif analysis was performed on (e) upregulated and (f) downregulated regions in monocytic cells relative to progenitor cells as described in panel d. Gray dots represent motifs that did not reach statistical significance (FDR < 0.05). **g.** Venn diagram displaying the overlap of significantly dysregulated transcription factors identified from single-cell RNA-seq analysis in panel a compared to enriched motifs identified in panels e and f.

We next performed single-cell ATAC-seq to characterize chromatin accessibility dynamics following venetoclax treatment. Each cluster was assigned a likely cell type by evaluating predicted activity of lineage-defining transcription factors (Supplementary Figure 6a-g). To identify transcriptional programs affected by drug treatment, we performed differential peak calling analysis, comparing peaks in progenitor and monocyte clusters (Supplementary Figure 6h,i). Motif enrichment analysis of regions preferentially accessible in monocytic populations compared to progenitor populations at 72 hours identified 325 transcription factor motifs that varied by differentiation state and were potentially influenced by venetoclax (Figure 4d-f; Supplementary Table 7). To link chromatin accessibility with downstream impacts on gene expression, we examined overlap between these differentially accessible motifs and the 62 transcription factors with higher predicted activity, identifying 27 transcription factors in common (Figure 4g). We eliminated transcription factors whose activity decreased over pseudotime and used known involvement in myeloid differentiation to nominate seven transcription factors likely involved in venetoclax resistance: PU.1, BACH1, CEBPβ, JUNB, MAFB, KLF6, and MXD1.

### Targeted CRISPR screening nominates PU.1 as a therapeutic target for enhancing venetoclax sensitivity

To identify genetic vulnerabilities in stem-monocytic AML, we performed a targeted CRISPR knockout screen of the seven transcription factors identified from our single-cell analyses, plus *BCL2*, *BCL2L1*, and the safe-harbor locus *AAVS1* as controls. We electroporated OCI-AML8227 cells with guide RNAs targeting each factor and measured immunophenotypic and viability profiles following DMSO or venetoclax treatment (Figure 5a). This approach enabled us to directly assess how loss of each factor influenced both response to BCL2 inhibition and differentiation state. Only *SPI1* knockout increased sensitivity to venetoclax compared with *AAVS1* controls (Figure 5b). Loss of these factors had little effect on differentiation, with the notable exception of *SPI1*, knockout of which entirely halted differentiation beyond the CD38⁺ stage (Figure 5c-j; Supplementary Figure 7). These results suggest that *SPI1* loss maintains leukemic cells in an immature state, thereby increasing their sensitivity to BCL2 inhibition. Together, these data show that PU.1 is a key determinant of differentiation-associated venetoclax response and nominate PU.1 blockade as a potential mechanism to enhance venetoclax sensitivity in stem-monocytic AML.

**Figure 5:**
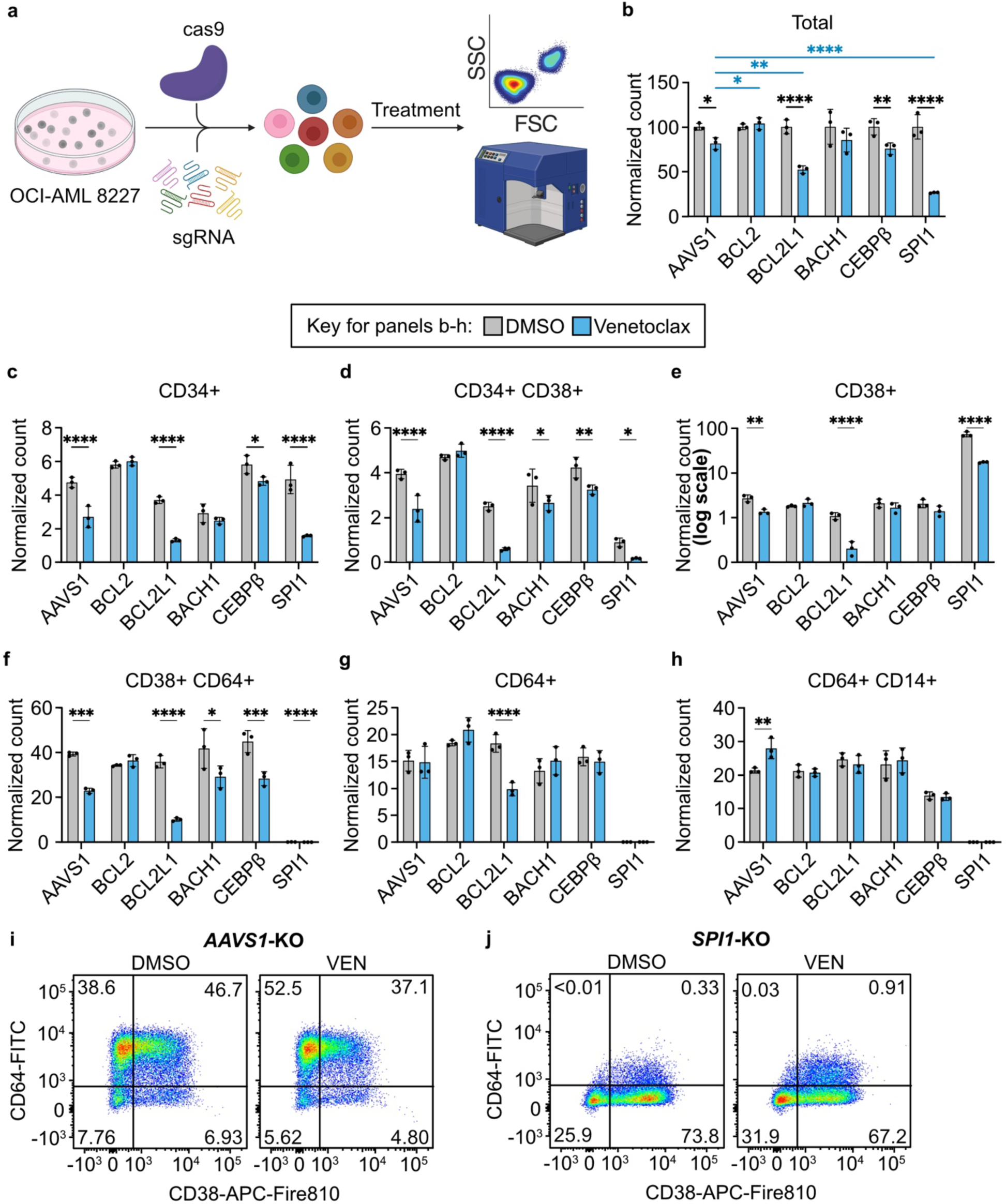
Targeted CRISPR screening nominates PU.1 as a therapeutic target for enhancing venetoclax sensitivity. **a.** Schematic of the targeted CRISPR screen. A stable Cas9-expressing OCI-AML8227 cell line was generated by electroporation and sorted for GFP expression. Cells were transduced with three guide RNAs per transcription factor target by electroporation, then cultured in triplicate with DMSO or 1 μM venetoclax for 72 hours before flow cytometry analysis of CD34, CD38, CD64, and CD14 surface expression. Image was created with BioRender. **b.** Live cell counts determined by forward/side scatter gating, excluding DAPI-stained cells. Counts were normalized to the average of their respective DMSO-treated controls. Blue lines and asterisks indicate comparisons with the safe-harbor locus *AAVS1* (control) whereas black lines and asterisks indicate intra-sample comparisons. Significance was evaluated using ordinary two-way ANOVA followed by Holm-Šidák post-test correction. **c–h.** Live cell counts for each immunophenotypic population, normalized as described in panel b. Significance was evaluated using ordinary two-way ANOVA followed by Holm-Šidák post-test correction. **i–j.** Representative flow cytometry plots showing CD64-FITC and CD38-APC-Fire surface expression in OCI-AML8227 cells transduced with guide RNAs targeting *AAVS1* or *SPI1* following treatment with DMSO or 1 μM venetoclax. ns = not significant; * = p < 0.05; ** = p < 0.01; *** = p < 0.001; **** = p < 0.0001.

### Pharmacologic PU.1 inhibition synergizes with venetoclax in preclinical models and primary patient samples

To improve the translatability of our targeted CRISPR knockout experiments, we evaluated pharmacologic strategies to inhibit PU.1. We identified DB2313, a small molecule that disrupts PU.1 binding and downregulates canonical PU.1 transcriptional targets, as a candidate agent^43,44^. OCI-AML8227 cells demonstrated sensitivity to DB2313 monotherapy with an IC50 of 5.9 μM (Figure 6a). Combined treatment with venetoclax and DB2313 revealed synergy at approximately 5 μM DB2313 and 1 μM venetoclax (Figure 6b). Immunophenotypic profiling revealed that even at low concentrations, the combination preferentially depleted the most immature subpopulations (Figure 6c-e; Supplementary Figure 8a-c). At higher concentrations associated with synergy, DB2313 produced more pronounced effects, resulting in near-complete depletion of CD34⁺ cells (Figure 6f-h; Supplementary Figure 8d-f). Consistent with *SPI1* knockout, DB2313 monotherapy at both concentrations induced differentiation arrest at the CD38⁺ stage, as evidenced by expansion of this population relative to DMSO controls. These findings demonstrate that pharmacologic PU.1 inhibition recapitulates the genetic loss-of-function phenotype and produces synergistic effects with venetoclax in stem-monocytic AML.

**Figure 6:**
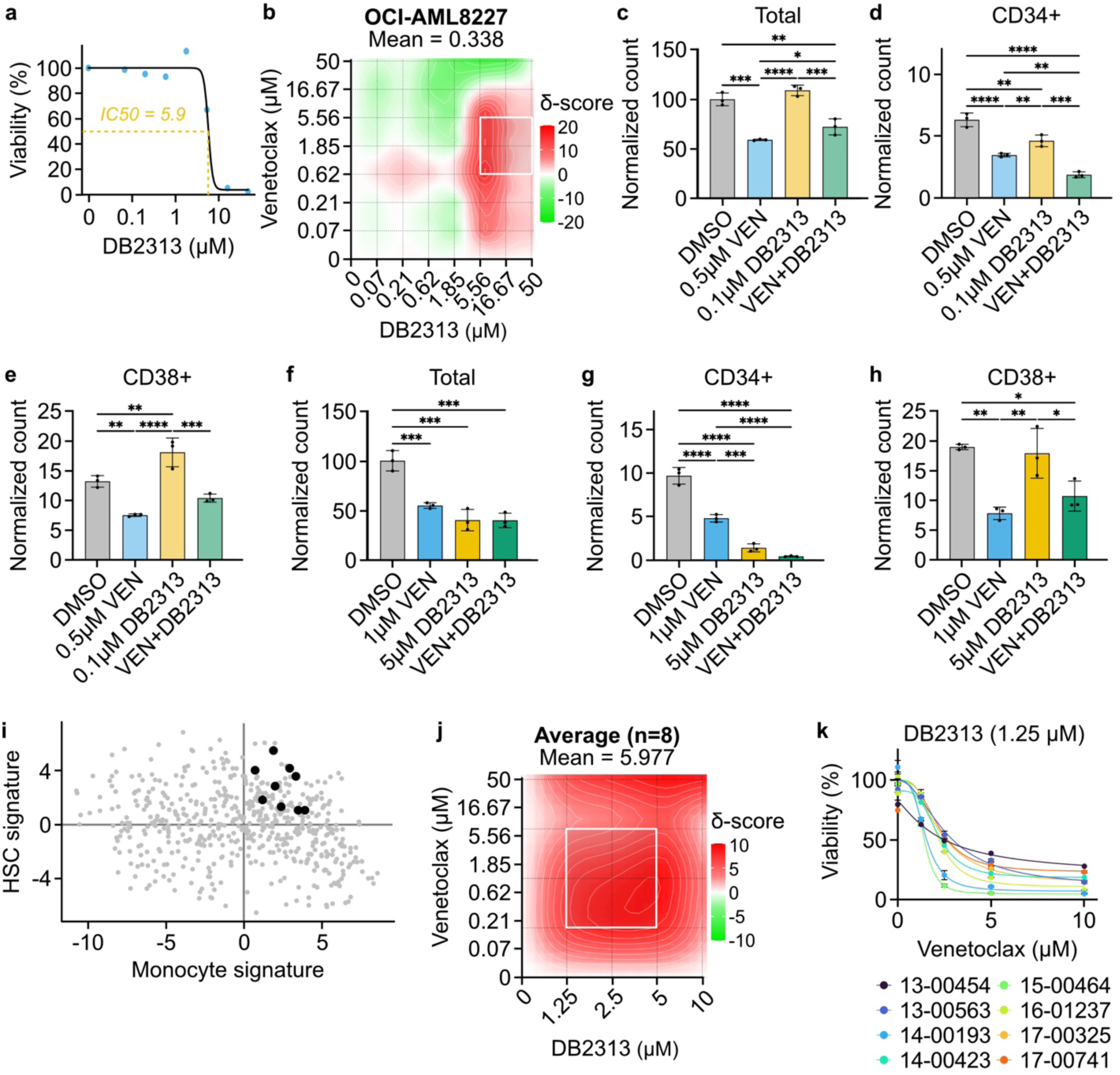
Pharmacologic PU.1 inhibition synergizes with venetoclax in preclinical models and primary patient samples. **a.** OCI-AML8227 cells were cultured in triplicate for 72 hours along an 8-point dose curve of DB2313. Cell viability was assessed by CellTiter Aqueous colorimetric assay. **b.** OCI-AML8227 cells were treated in triplicate with an 8×8 dose matrix of DB2313 and venetoclax for 72 hours prior to viability assessment by CellTiter Aqueous colorimetric assay. Zero interaction potency (ZIP) synergy scores were calculated on the average values for each drug dose. The white box indicates the DB2313 and venetoclax concentrations corresponding to maximal synergy. **c–e.** Live cell counts in OCI-AML8227 cells following 72 hours of treatment with 0.1 μM venetoclax, 0.5 μM DB2313, both drugs in combination, or an equivalent volume of DMSO. Live cell counts were determined by forward/side scatter gating and exclusion of DAPI-stained cells, then normalized to DMSO-treated controls. Cells were analyzed by flow cytometry for CD34, CD38, CD64, and CD14 surface expression. Quantification of live cell counts for the remaining cell surface markers is shown in Supplementary Figure 8. Significance was evaluated using ordinary two-way ANOVA followed by Holm-Šidák post-test correction. **f–h.** Live cell counts in OCI-AML8227 cells following 72 hours of treatment with 1 μM venetoclax, 5 μM DB2313, both drugs in combination, or an equivalent volume of DMSO. Analysis was performed as described in panels c–e. **i.** Transcriptional signatures of nine primary AML samples selected for drug sensitivity evaluation are shown. One sample was excluded from downstream analyses due to widespread cell death (18-00105). **j.** Primary AML blasts from eight patients with stem-monocytic AML were cultured in triplicate for 72 hours along a 7-point dose curve with venetoclax, DB2313, or equimolar amounts of the drug combination. Viability was assessed using the Guava/EMD Millipore platform after a short incubation with Guava Nexin Reagent (Annexin V–PE + 7-AAD). ZIP synergy scores were calculated from averaged viability data across replicates for each drug dose in primary AML blasts shown in panel i. The white box indicates the DB2313 and venetoclax concentrations corresponding to maximal synergy. **k.** Venetoclax dose-response curves for each patient at a fixed dose of 1.25 μM DB2313, which corresponds to maximal synergy in panel j. ns = not significant; * = p < 0.05; ** = p < 0.01; *** = p < 0.001; **** = p < 0.0001.

To assess the clinical relevance of the DB2313 and venetoclax combination, we validated this approach in eight primary stem-monocytic patient samples (Figure 6i). DB2313 enhanced venetoclax activity in nearly every sample tested, and the combination demonstrated statistically significant synergy across the entire patient cohort (Figure 6j,k; Supplementary Figure 9). Together, these data reveal that concurrent BCL2 and PU.1 inhibition enhances stem-monocytic leukemic cell death, likely through induction of a differentiation block that maintains cells in a venetoclax-sensitive immature state.

To assess the broader applicability of this combination strategy, we evaluated DB2313 and venetoclax across a diverse panel of AML cell lines representing distinct genetic and phenotypic subtypes (Supplementary Figure 10). Significant synergy was observed in nearly all cell lines tested, with the exception of OCI-AML3. The breadth of synergistic responses across this genetically and phenotypically diverse panel suggests therapeutic benefit extends beyond stem-monocytic AML to encompass multiple AML subtypes.

## DISCUSSION

In this work, we describe stem-monocytic AML, a transcriptionally distinct patient population exhibiting features of both HSC-like and monocyte-like leukemia. Analysis of *ex vivo* drug-sensitivity data from a large patient cohort showed that stem-monocytic AML is as resistant to venetoclax as monocytic AML. We were unable to identify consistent genetic or immunophenotypic patterns in this population, emphasizing the limitations of current diagnostic methods and the importance of transcriptional differentiation state as a functional biomarker. Using a leukemia stem cell model this transcriptional subtype, we found that venetoclax treatment preferentially depleted immature blasts while enriching for monocytic leukemic populations marked by increased PU.1 activity. CRISPR-mediated PU.1 knockout induced differentiation arrest and enhanced venetoclax sensitivity. Similarly, pharmacologic disruption of PU.1-DNA binding with the small molecule DB2313 increased venetoclax efficacy in both AML model systems and primary stem-monocytic AML patient samples. Together, these findings identify stem-monocytic AML as a transcriptionally and functionally distinct subtype and nominate combined PU.1 and BCL2 inhibition as a rational therapeutic strategy for this patient population.

The absence of identifiable genetic or immunophenotypic patterns in stem-monocytic AML suggests that conventional diagnostic approaches, including mutational profiling and flow cytometry, may be unable to reliably detect this subtype despite its representation among a substantial fraction of patients. This limitation highlights that transcriptional differentiation state captures biological features not discernible through standard clinical testing. Notably, the Beat AML dataset encodes only binary presence of cell surface markers^8,45^. Incorporating quantitative assessment of population proportions may improve detection of these cases. Moreover, larger patient cohorts will be required to determine whether recurrent genetic alterations underlie this transcriptional phenotype. Collectively, these observations establish stem-monocytic AML as a clinically relevant population and suggest that transcriptional differentiation profiling could serve as a critical biomarker for patient identification^46^.

The PU.1 small-molecule inhibitor used in this study consistently enhanced venetoclax efficacy across model systems and primary patient samples. However, this compound acts indirectly by redistributing PU.1 binding at a subset of its genomic targets^47^. Alternative strategies capable of achieving more direct or comprehensive PU.1 inhibition warrant further evaluation in this population. Potential approaches include targeted protein degradation via PROTACs or transcript suppression through antisense oligonucleotides^48^. Such agents may offer greater potency and improved selectivity across PU.1-regulated transcriptional networks, potentially broadening therapeutic applicability in this patient population.

These findings establish a rational therapeutic combination for stem-monocytic AML that warrants clinical investigation. Determining whether transient or partial PU.1 inhibition affects hematologic recovery will be essential for clinical development. Particular attention should be given to the hematopoietic consequences of PU.1 inhibition, given the high frequency of leukopenia in patients receiving venetoclax-based regimens^43,44^. Prospective trials including this patient population will clarify whether PU.1 targeting can be safely combined with BCL2 inhibition and provide mechanistic insight into differentiation state-dependent therapeutic responses.

## CONCLUSION

Stem-monocytic AML is a transcriptionally and functionally distinct subtype that is not identifiable by conventional clinical testing. This population exhibits limited venetoclax response mediated by myeloid differentiation programs regulated by PU.1. Combined PU.1 and BCL2 inhibition enhances venetoclax sensitivity in both model systems and patient samples, establishing a potential strategy for improving therapeutic response in this population. Future clinical investigations that integrate transcriptional profiling with functional drug-sensitivity testing will be critical for validating this therapeutic approach and refining the classification of differentiation-based AML subtypes.

## AUTHORS’ CONTRIBUTIONS

**WM Yashar**: conceptualization, formal analysis, funding acquisition, investigation, methodology, software, validation, visualization, writing – original draft, writing – review & editing, final manuscript approval. **IV Pacentine**: formal analysis, investigation, methodology, software, validation, visualization, writing – original draft, writing – review & editing, final manuscript approval. **A Taherinasab**: investigation, methodology, validation, final manuscript approval. **T Nguyen**: formal analysis, investigation, final manuscript approval. **S Worme**: formal analysis, investigation, final manuscript approval. **M Tsuchiya**: investigation, methodology, validation, final manuscript approval. **T Lusardi**: investigation, supervision, final manuscript approval. **C Posso**: data curation, formal analysis, investigation, software, visualization, final manuscript approval. **P Piehowski**: data curation, formal analysis, investigation, software, supervision, final manuscript approval. **S Gosline**: investigation, supervision, final manuscript approval. **GG Yardimci**: investigation, methodology, resources, software, supervision, final manuscript approval. **AC Adey**: investigation, methodology, resources, software, supervision, final manuscript approval. **JE Maxon**: conceptualization, investigation, resources, supervision, writing – review & editing, final manuscript approval. **TP Braun**: conceptualization, formal analysis, funding acquisition, investigation, methodology, resources, supervision, writing – review & editing, final manuscript approval.

## ACKNOWLEDGEMENTS

We would like to thank all of the Beat AML patients for their generous donation of time and biospecimens in support of this research. We are also deeply grateful to the physicians, nurses, clinical coordinators, and support staff who cared for these patients and continue to provide exceptional care for individuals with leukemia. Their dedication makes this work possible. We appreciate the OHSU ExaCloud Cluster Computational Resource and the Advanced Computing Center for their assistance. The OCI-AML8227 cells were generously provided by Dr. John Dick (University of Toronto). A special thank you to Madison Hall for optimizing the protocol for and maintaining our lab supply of 5637 cell media, a constituent of MUTZ3 media. This work was funded by the National Institutes of Health (1F30CA278500-01A1 to WM Yashar, K08 CA245224 and 1R01 CA282133-01 to TP Braun), the American Cancer Society (P30 CA016672-40 to TP Braun), and the American Society of Hematology Research Restart and Scholar Awards to TP Braun.

## SUPPLEMENTARY FIGURE LEGENDS

**Supplementary Figure 1:**
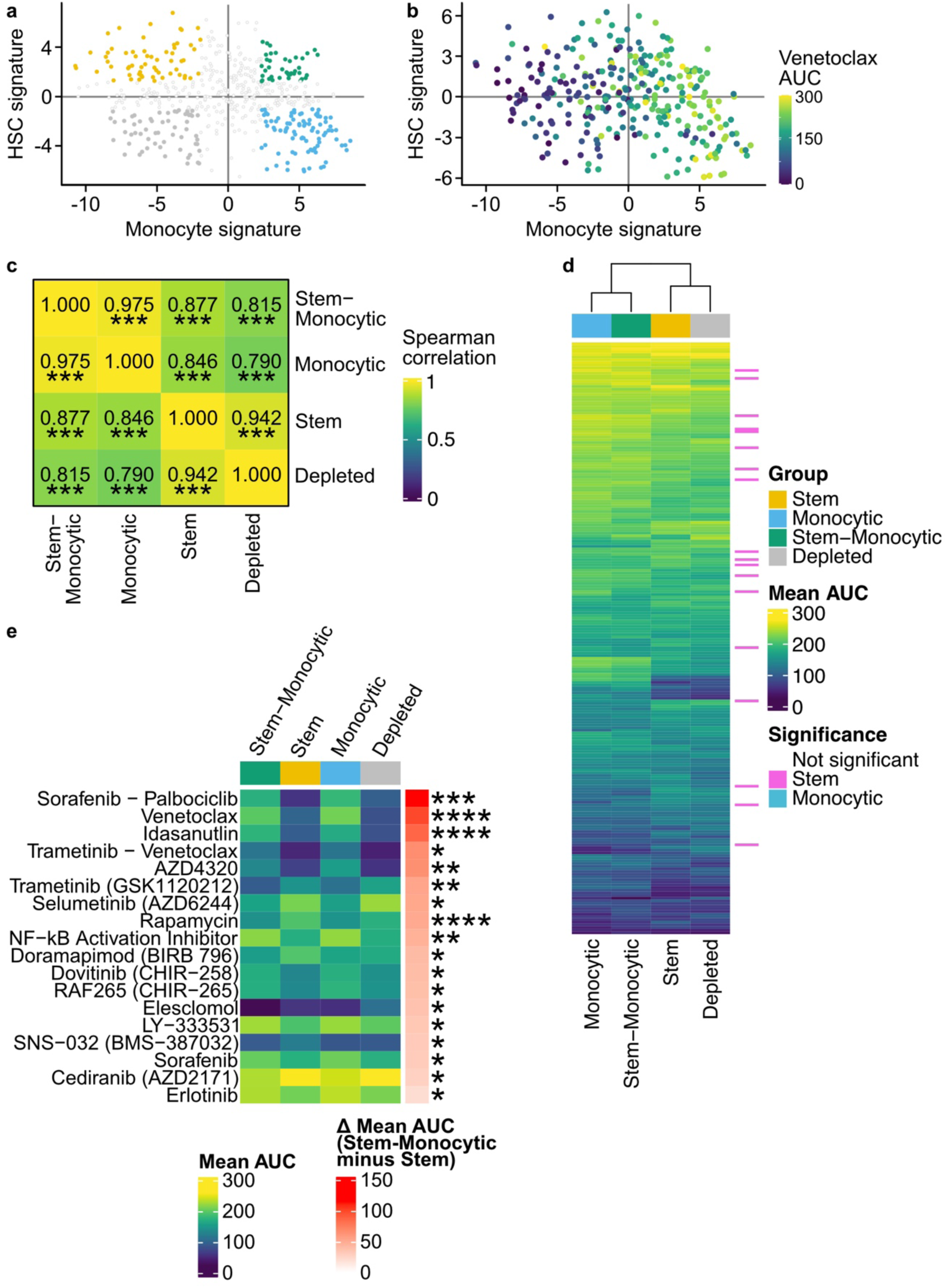
Stem-monocytic AMLs exhibit strong concordance with monocytic AMLs in their drug-response signatures. **a.** Cell type transcriptional signatures were calculated using previously described methods for 652 patients in the BeatAML cohort^8^. The first principal component of the hematopoietic stem cell (HSC) and monocyte signatures is visualized, where each point represents a single patient as shown in Figure 1a. Patients were stratified based on HSC and monocyte transcriptional signatures. Colored dots indicate patients within the top 33% percentile of HSC and monocytic signatures (total n = 289, stem n = 73, stem-monocytic n = 52, monocytic n = 99, depleted n = 65). Gray dots represent patients that did not reach this threshold. The four colors correspond to the transcriptional subtypes: stem, depleted, monocytic, and stem-monocytic. **b.** Primary AML blasts from 652 patient samples were cultured in triplicate for 72 hours with a panel of small-molecule inhibitors across 7-point dose curves. Dose-response curves for all the drugs were fit using probit models and area under the curve (AUC) values were calculated for each inhibitor within each transcriptional subtype. Cell viability was assessed by CellTiter Aqueous colorimetric assay. AUC values from the dose-response curves are overlaid onto the HSC and monocyte signatures for each patient as shown in Figure 1a. **c.** Primary AML blasts from 652 patient samples representing the top 33% of each transcriptional subtype (as shown in panel a) were cultured in triplicate for 72 hours with a panel of small-molecule inhibitors across 7-point dose curves. Dose-response curves for all the drugs were fit using probit models and AUC values were calculated for each inhibitor within each transcriptional subtype. Median inhibitor responses across subtypes were linearly correlated, with significance determined by two-sided Spearman tests. **d, e.** Annotated heatmap of scaled inhibitor responses ordered by hierarchical clustering. Row annotations indicate inhibitors with significant differences in response between stem-monocytic and stem AML, and between stem-monocytic and monocytic AML, as determined by p-value < 0.05 from multiple t-tests adjusted with Benjamini-Hochberg correction. Only inhibitors with at least 10 samples per group were included in the analysis (total n = 225, stem n = 51, stem-monocytic n = 41, monocytic n = 81, depleted n = 52). ns = not significant; * = p < 0.05; ** = p < 0.01; *** = p < 0.001; **** = p < 0.0001.

**Supplementary Figure 2:**
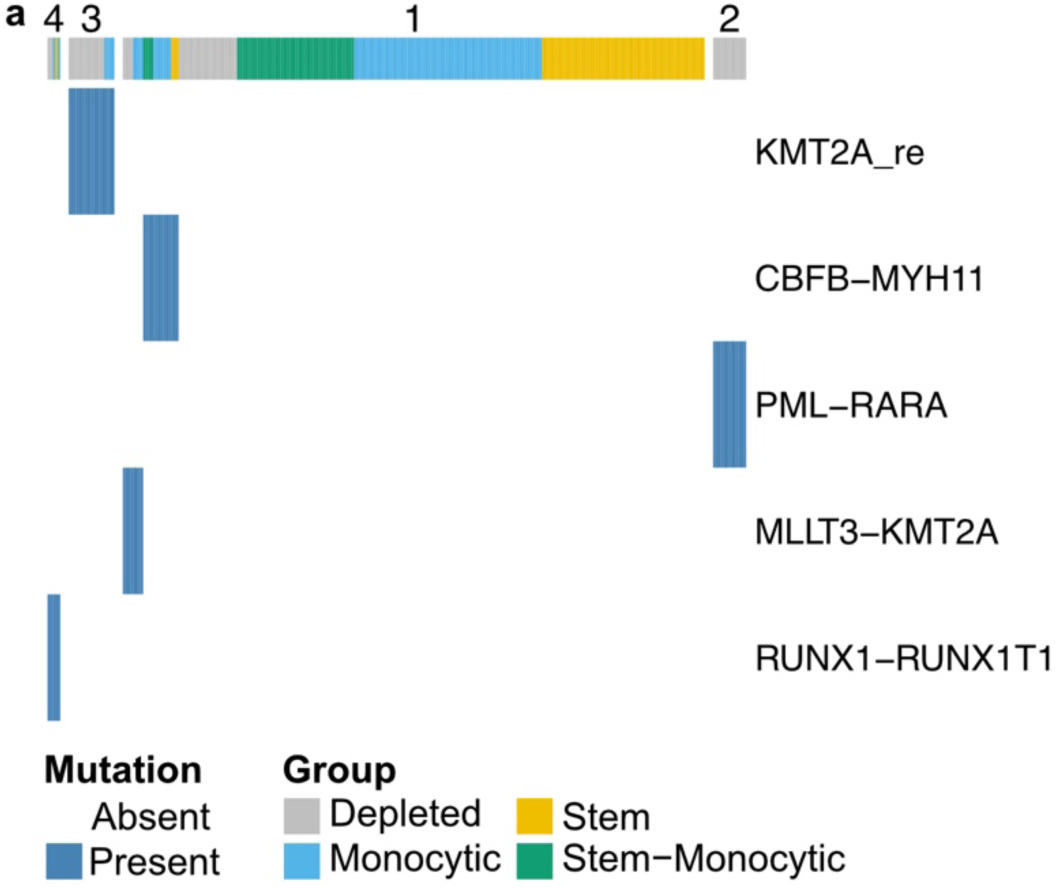
Stem-monocytic AML is not characterized by common gene fusions. **a.** Gene fusions for each patient were identified using conventional clinical methods, including karyotyping, FISH, and RT-PCR. Cell type transcriptional signatures were calculated using previously described methods for 652 patients in the BeatAML cohortt^8^. Patients within the top 33% of each transcriptional subtype were selected for analysis. Each patient is color-coded according to subtype.

**Supplementary Figure 3:**
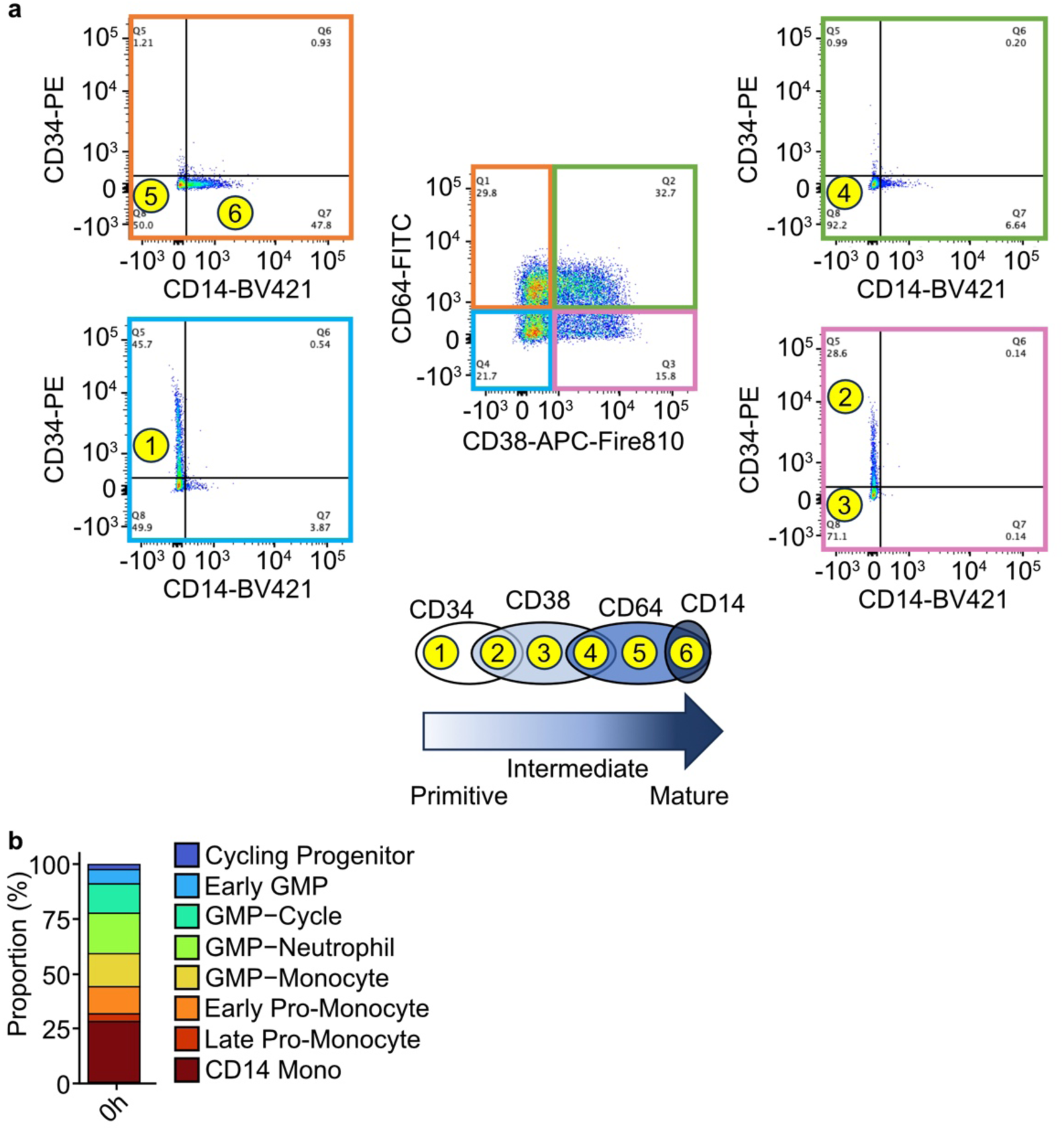
Common surface markers associated with myeloid differentiation were used to characterize the differentiation state of OCI-AML8227 cells. **a.** Gating strategy used to quantify proportions of cell types. The six main subpopulations of OCI-AML8227 cells are labeled in yellow circles, representing a continuum from immature to intermediate as shown in the bottom panel. **b.** Quantification of the proportions of cell types in scRNA-seq data from untreated OCI-AML8227 cells.

**Supplementary Figure 4:**
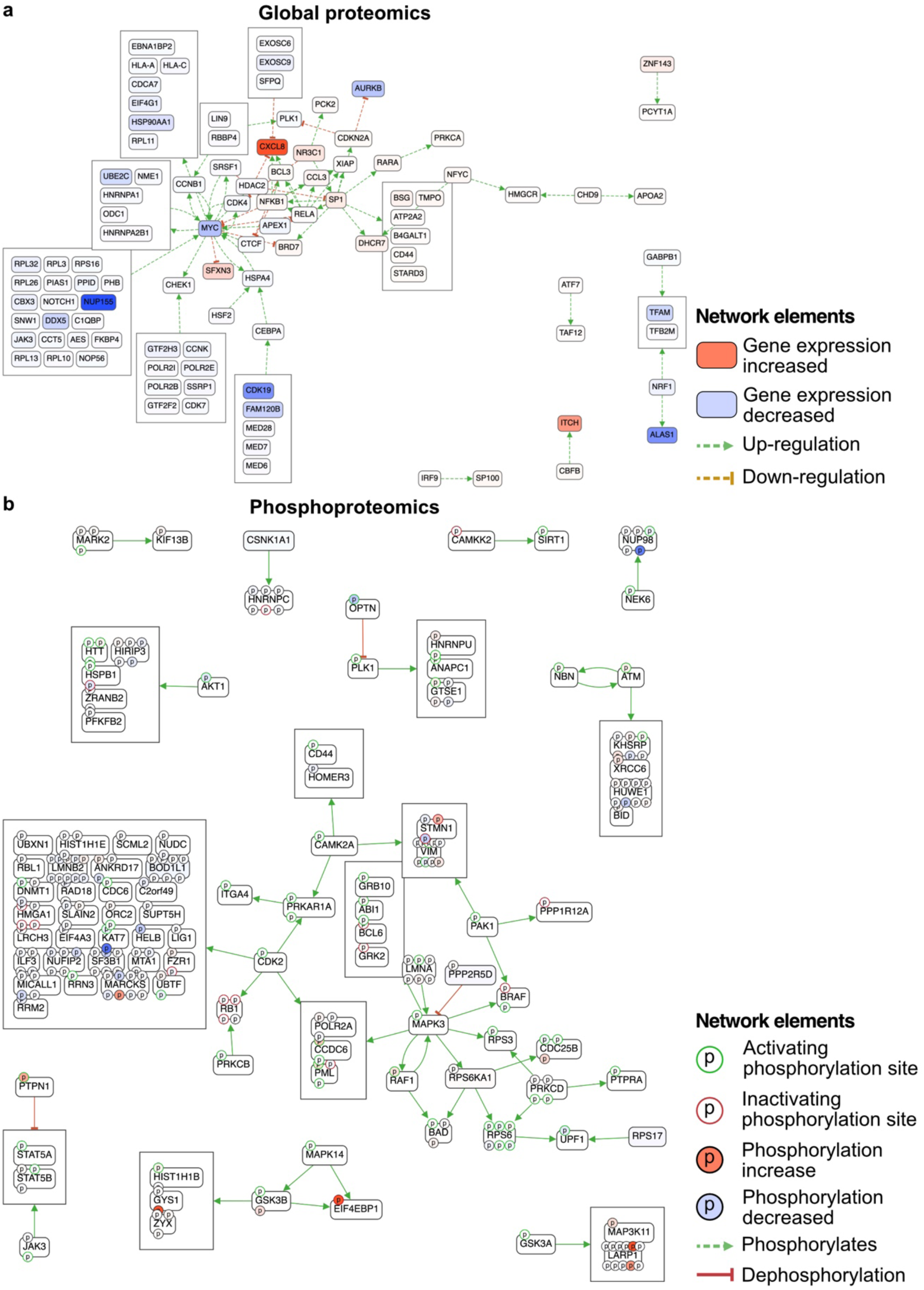
Proteomics reveal that BCL2 inhibition disrupts central cell cycle and apoptotic regulators in immature leukemic cells. **a, b.** OCI-AML8227 cells were immunomagnetically fractionated by CD34 expression and cultured in triplicate for 30 minutes or 6 hours following treatment with 1 μM venetoclax or an equivalent volume of DMSO. Proteins were extracted from cell pellets and either separated by liquid chromatography and analyzed by tandem mass spectrometry for protein identification and quantification or enriched for phosphorylated species prior to LC–MS/MS analysis to profile phosphorylation-dependent signaling. Relative (a) global protein and (b) phosphoprotein abundance at 6 hours following venetoclax treatment in CD34-enriched and CD34-depleted populations were analyzed using proteomic network analysis.

**Supplementary Figure 5:**
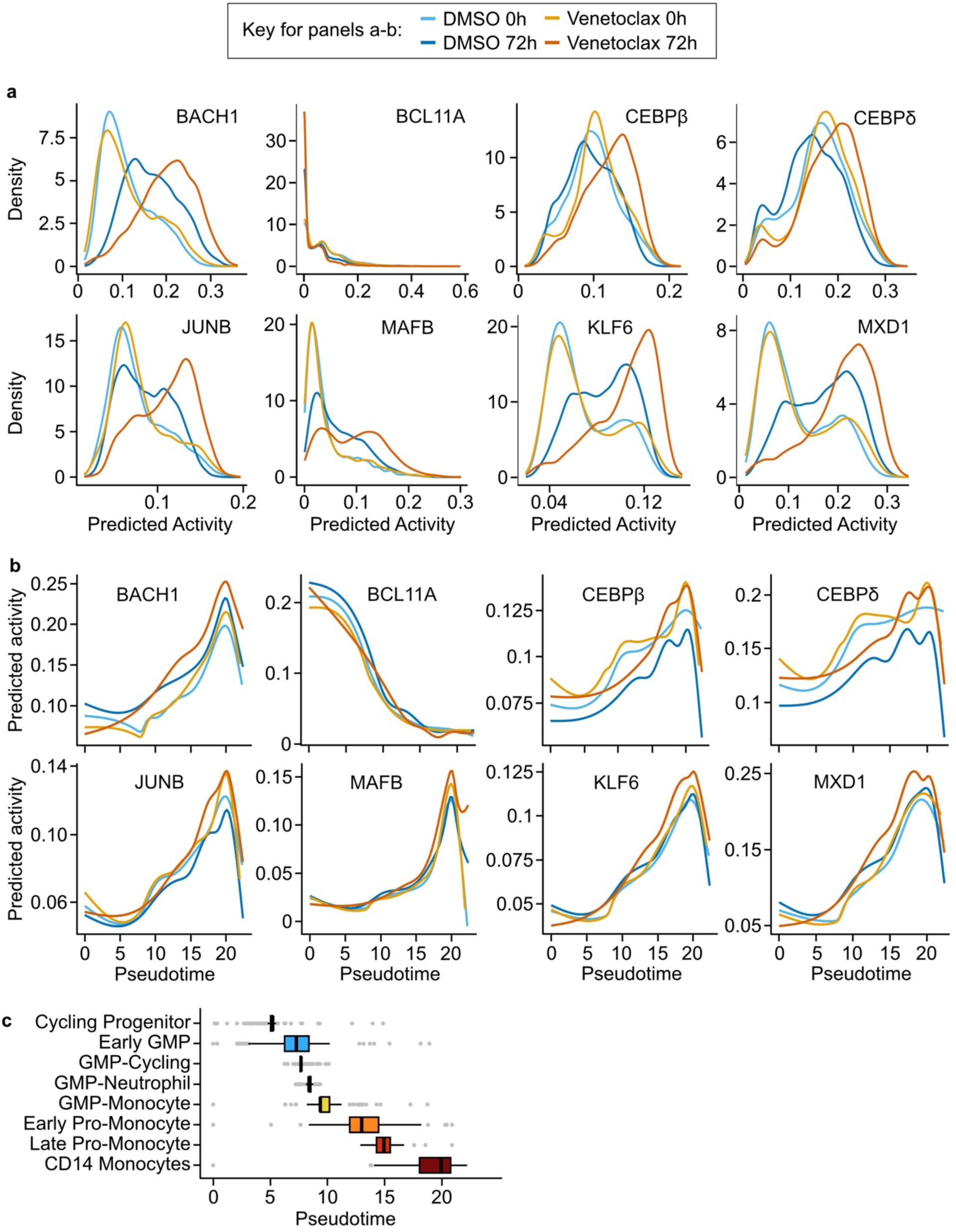
Putative transcription factors associated with venetoclax resistance are also associated with myeloid differentiation. **a.** Predicted transcription factor activity was evaluated in scRNA-seq data presented in Figure 3. Distributions of predicted transcription factor activity for different transcription factors in each drug condition. **b.** Predicted activity from panel a plotted along the myeloid differentiation trajectory (pseudotime) in each drug condition. **c.** Boxplot of the pseudotime per cell type in OCI-AML8227 cells.

**Supplementary Figure 6:**
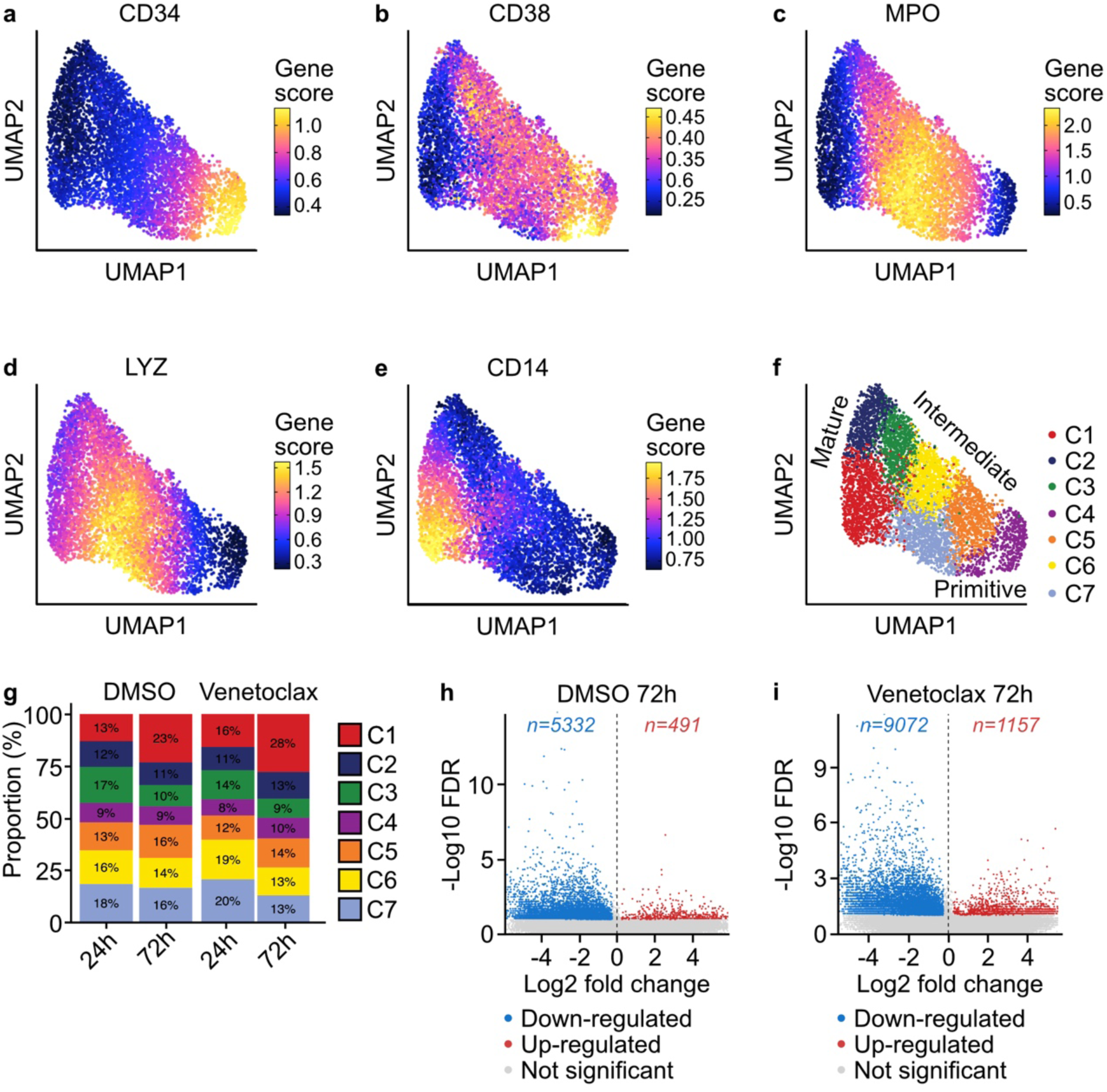
Single-cell chromatin accessibility profiles nominate regions enriched in monocytic cells compared to progenitors. **a–e.** OCI-AML8227 cells were cultured with DMSO or 1 μM venetoclax and processed at either 0 or 72 hours for single-cell ATAC-seq. UMAP dimensional reduction and projection of single-cell chromatin accessibility profiles from OCI-AML8227 cells in four conditions. Predicted activity of various transcription factors associated with distinct stages in myeloid differentiation. **f.** Unsupervised clustering was performed on regions of open chromatin accessibility. Clusters were broadly categorized as mature, intermediate, or immature based on predicted activity of multiple transcription factors involved in myeloid differentiation as shown in panels a–e. **g.** Stacked bar plots displaying the proportions of cells belonging to each cluster described in panel f for each sample. **h, i.** Peak calling identified regions of chromatin accessibility in each cluster. Volcano plots display regions upregulated and downregulated in monocytic clusters compared to progenitor clusters in cells treated with (h) DMSO and (i) venetoclax at 72 hours. Gray dots represent peaks that did not reach statistical significance (FDR < 0.05).

**Supplementary Figure 7:**
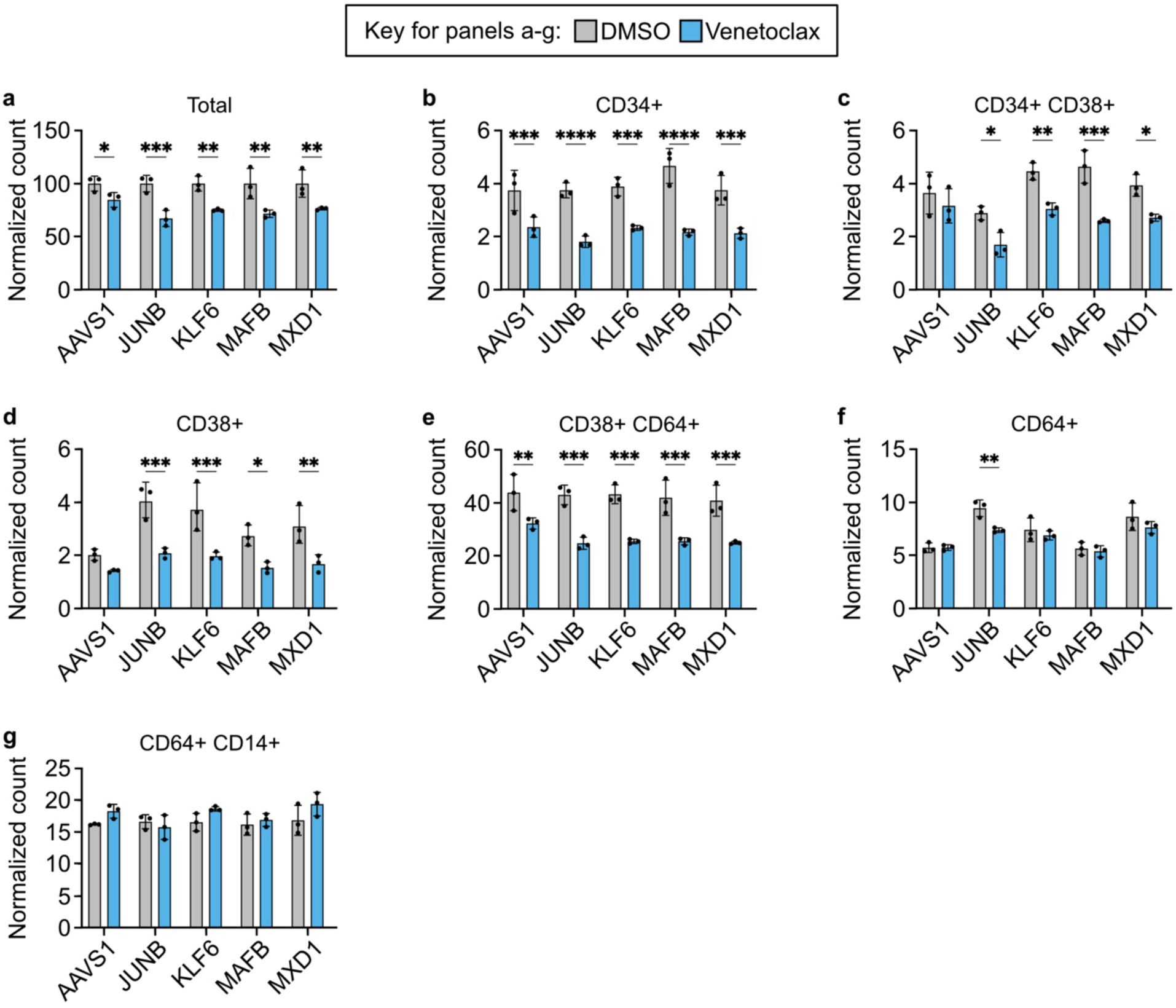
CRISPR knockdown of transcription factors with little effect on differentiation. **a.** Cells were transduced with three guide RNAs per transcription factor target by electroporation, then cultured in triplicate with DMSO or 1 μM venetoclax for 72 hours before flow cytometry analysis of CD34, CD38, CD64, and CD14 surface expression as described in Figure 5a. Quantification of total cell counts in additional knockout models of OCI-AML8227 cells. Total cell counts were normalized to the average of their respective DMSO-treated controls. Significance was evaluated using ordinary two-way ANOVA followed by Holm-Šidák post-test correction. Sensitivity to venetoclax was unchanged by the experimental knockouts compared to control *AAVS1*. **b–g.** Live cell counts for each immunophenotypic population were calculated by multiplying their proportions of live single cells from flow cytometric analysis by their normalized values in panel a. Significance was evaluated using ordinary two-way ANOVA followed by Holm-Šidák post-test correction. Few changes were observed in knockouts relative to control. ns = not significant; * = p < 0.05; ** = p < 0.01; *** = p < 0.001; **** = p < 0.0001.

**Supplementary Figure 8:**
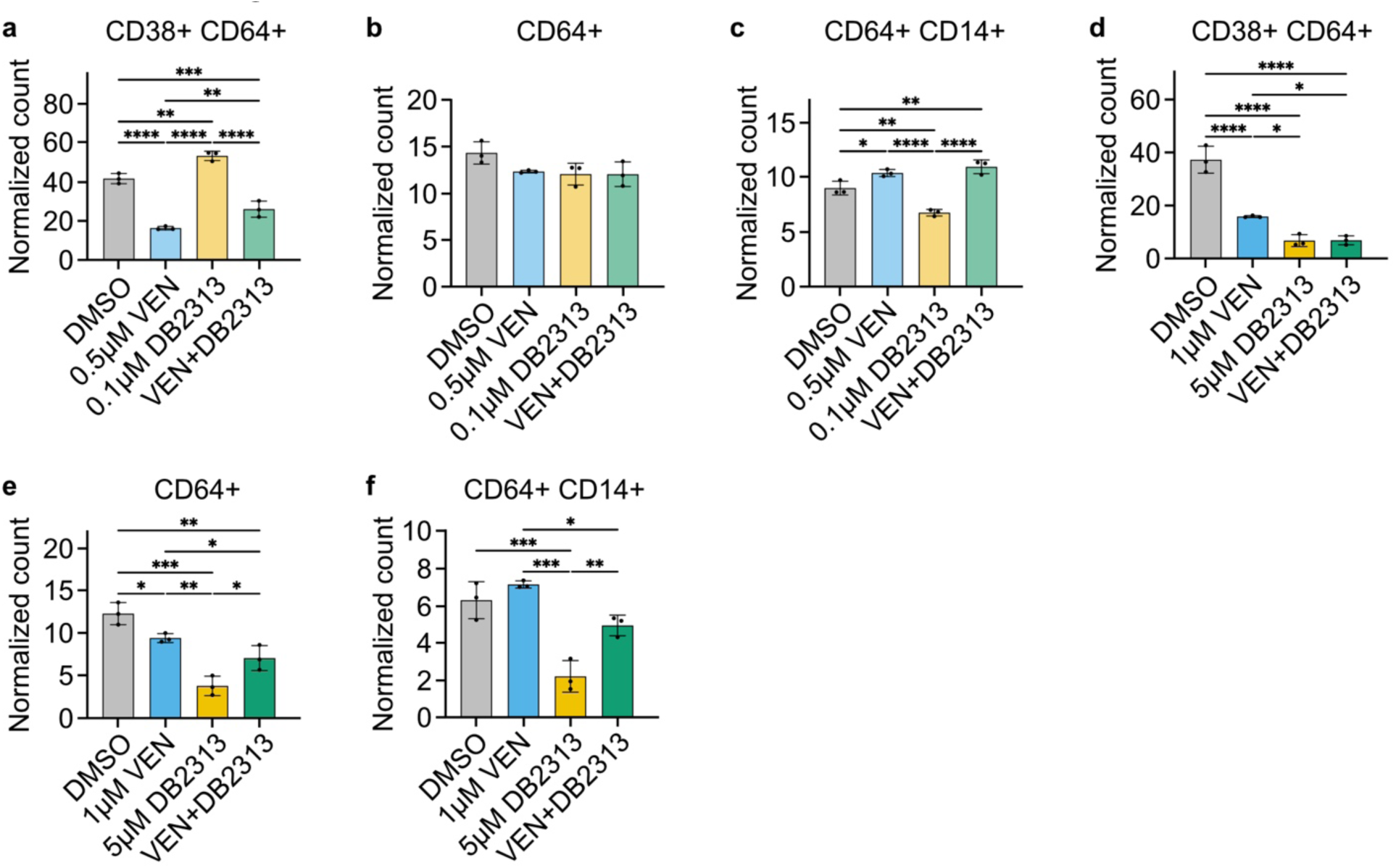
Combined BCL2 and PU.1 inhibition preferentially depletes the most immature subpopulations at low and high concentrations. **a–c.** Live cell counts in OCI-AML8227 cells following 72 hours of treatment with 0.1 μM venetoclax, 0.5 μM DB2313, both drugs in combination, or an equivalent volume of DMSO. Analysis was performed as described in Figure 6c–e. **d–f.** Live cell counts in OCI-AML8227 cells following 72 hours of treatment with 1 μM venetoclax, 5 μM DB2313, both drugs in combination, or an equivalent volume of DMSO. Analysis was performed as described in Figure 6c–e. ns = not significant; * = p < 0.05; ** = p < 0.01; *** = p < 0.001; **** = p < 0.0001.

**Supplementary Figure 9:**
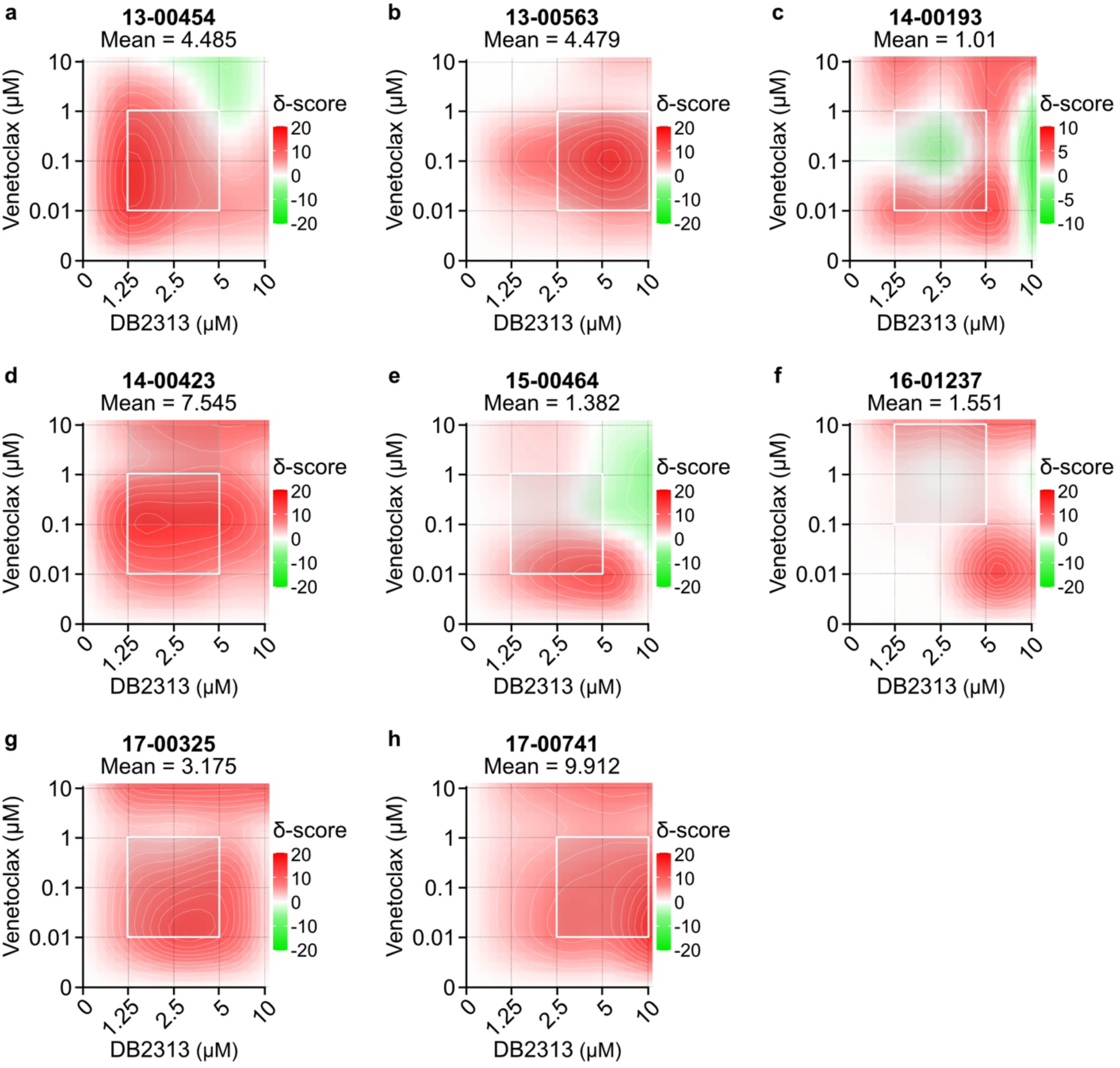
Concurrent BCL2 and PU.1 inhibition enhances stem-monocytic leukemic cell death in primary patient samples. **a–h.** Primary AML blasts from nine patients with stem-monocytic AML were cultured in triplicate for 72 hours along a 7-point dose curve with venetoclax, DB2313, or equimolar amounts of the drug combination. Viability was assessed using the Guava/EMD Millipore platform after a short incubation with Guava Nexin Reagent (Annexin V–PE + 7-AAD). One sample was excluded from downstream analyses due to widespread cell death (18-00105). ZIP synergy scores were calculated from averaged viability data across replicates for each drug dose in primary AML blasts. The white boxes indicate the DB2313 and venetoclax concentrations corresponding to maximal synergy.

**Supplementary Figure 10:**
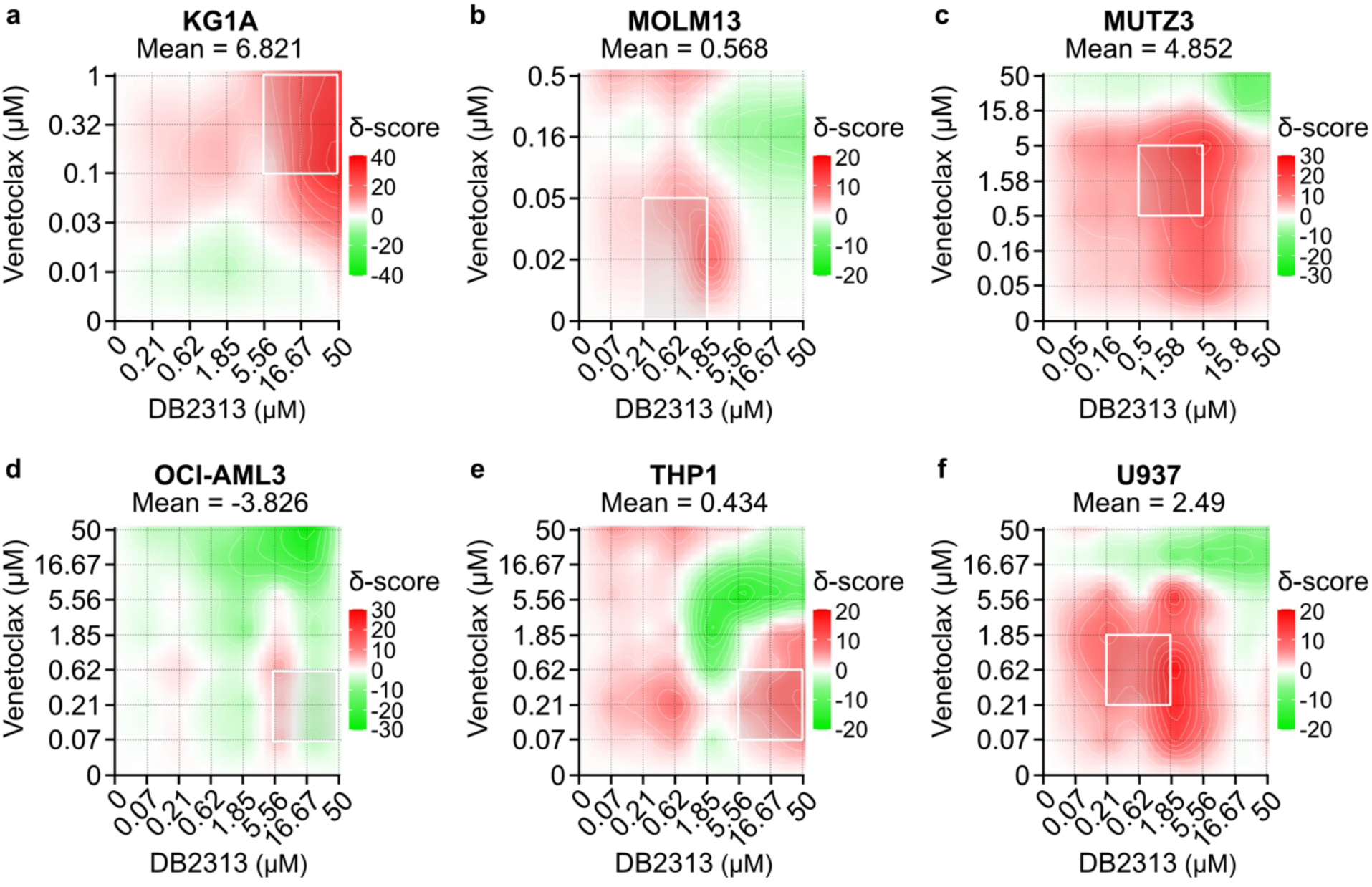
Pharmacologic inhibition of PU.1 with DB2313 synergizes with venetoclax in several genetically and phenotypically distinct AML models. **a–f.** Each cell line was treated in triplicate with an 8×8 dose matrix of DB2313 and venetoclax (7×7 for KG1A cells) for 72 hours prior to viability assessment by CellTiter Aqueous colorimetric assay. ZIP synergy scores were calculated from averaged viability data across replicates for each drug dose. The white boxes indicate the DB2313 and venetoclax concentrations corresponding to maximal synergy.

